# Genetics of continuous colour variation in a pair of sympatric sulphur butterflies

**DOI:** 10.1101/2023.02.03.526907

**Authors:** Joseph J. Hanly, Caroline M. Francescutti, Ling S. Loh, Olaf B. W. H. Corning, Derek J. Long, Marshall A Nakatani, Adam H. Porter, Arnaud Martin

## Abstract

Continuous colour polymorphisms can serve as a tractable model for the genetic and developmental architecture of traits, but identification of the causative genetic loci is complex due to the number of individuals needed, and the challenges of scoring continuously varying traits. Here we investigated continuous colour variation in *Colias eurytheme* and *C. philodice*, two sister species of sulphur butterflies that hybridise in sympatry. Using Quantitative Trait Locus (QTL) analysis of 483 individuals from interspecific crosses and an high-throughput method of colour quantification, we found that two interacting large effect loci explain around 70% of the heritable variation in orange-to-yellow chromaticity. Knockouts of *red Malphighian tubules* (*red*), a candidate gene at the primary QTL likely involved in endosomal maturation, resulted in depigmented wing scales showing disorganised pterin granules. The Z sex chromosome contains a large secondary colour QTL that includes the transcription factor *bric-a-brac* (*bab*), which we show can act as a modulator of orange pigmentation in addition to its previously-described role in specifying UV-iridescence. We also describe the QTL architecture of other continuously varying traits, and that wing size maps to the Z chromosome, supporting a Large-X effect model where the genetic control of species-defining traits is enriched on sex chromosomes. This study sheds light on the genetic architecture of a continuously varying trait, and illustrates the power of using automated measurement to score phenotypes that are not always conspicuous to the human eye.

**Foreword:** The colour phenotypes in this article involve nuanced gradations of yellow and orange that may be difficult to perceive for people who are colour vision deficient. Hue-shifted versions of all main figures are accessible online for dichromat readers (BioRxiv preprint: Supplementary Material).

## Introduction

Colour polymorphisms play key roles in adaptation and sexual selection in plants and animals (Orteu and Jiggins, 2020; Sapir et al., 2021; Wellenreuther et al., 2014), and provide a tractable system for the study of genotype-to-phenotype variation in natural populations (Orteu and Jiggins, 2020; San-Jose and Roulin, 2017). Phenotypic diversification and divergence in the colours of natural populations can vary discretely and be linked to one or few large effect loci – for example skin colour in wall lizards (Andrade et al., 2019), plumage colour in siskins and canaries (Gazda et al., 2020), or scale colours in *Heliconius* butterflies (Westerman et al., 2018) – while elsewhere it may vary continuously due to polygenic architectures consisting of many small effect loci – for example with human hair colour (Morgan et al., 2018), cichlid fish pigmentation (Albertson et al., 2014), or blue iridescence in butterflies (Brien et al., 2022, 2019). In the era of falling sequencing costs and big-data phenomics, study of continuous colour polymorphisms could provide fertile ground for improving our understanding of the genetic architecture of continuous traits.

A large number of studies have described the Mendelian genetics of colour variation in animals and plants. Various modes of colour production have been investigated, including the synthesis and deposition of melanin, carotenoid, psittacofulvin, and biliverdin pigments in vertebrates (Elkin et al., 2022); melanin, ommochrome, and pterin pigments in insects (Figon and Casas, 2019; Futahashi and Osanai-Futahashi, 2021); and the differentiation of iridescent structural features (Spiewak et al., 2018). These Mendelian colour polymorphisms have often been linked to a set of ‘hotspot’ pigmentation genes that were first described in model organisms, for example *MC1R* and *agouti* for melanin variation and *BCO2* for carotenoids (Elkin et al., 2022; Gazda et al., 2020).

More recently, studies in non-model organisms have identified new genetic mechanisms that underpin colour variation, independent of the need for a candidate gene approach. This is exemplified by the identification of recently-described enzymes Cyp2J19 and BDH1L in the carotenoid biosynthesis pathway, where a combination of genome wide association studies (GWAS), quantitative trait locus (QTL) mapping, and functional validation illustrated that these enzymes convert dietary carotenoids into red ketocarotenoids in birds (Toomey et al., 2022), and have likely acquired their role in body pigmentation through regulatory co-option from their function in eyes (Twyman et al., 2016). Continuing broad surveys of colour diversity are likely to uncover additional novel mechanisms for pigmentation across the tree of life.

Mendelian colour loci have made excellent systems for studying the genetic architecture of variation in natural populations, allowing for preliminary assessments of common classes of mutation; for example, spatial variation in pigmentation is found to be caused by *cis*-regulatory variation more often than coding variation (Elkin et al., 2022), and signalling molecules such as ligands and their receptors are recurring mutational targets of colour pattern evolution (Martin and Courtier-Orgogozo, 2017). While these findings inform our general understanding of the genetic architecture of other Mendelian traits, most observable traits in wild populations are non-mendelian. Robust studies of continuous colour variation are also needed to reflect the predominance of more complex traits that are pervasive in nature (Rockman, 2012). Continuous colour polymorphisms provide an opportunity to observe how spectra of allele effect sizes can vary in different evolutionary contexts. It is common in the wild (Roulin, 2016; Svensson and Wong, 2011), and can evolve rapidly in response to environmental perturbation (Burraco and Orizaola, 2022). Theoretical models predict that a combination of large effect and small effect loci will evolve in response to selection (Orr, 1998). A study of *Drosophila melanogaster* abdominal melanisation found that 3 large effect loci explain one third of colour heritability, with as many as 17 additional smaller-effect variants contributing (Dembeck et al., 2015): a similar architecture seems to be at play among two other, recently-diverged *Drosophila* species (Carbone et al., 2005; Yeh and True, 2014). Similarly in zebra finches, beak redness depends on environmental interactions, but has a heritable component that is partially explained by 4 linkage groups (Schielzeth et al., 2012; Slate, 2013). In contrast, in the polygenic white head patch of *Ficedula* flycatchers, no large-effect loci contribute to colour, meaning the phenotype is inherited via many small-effect loci (Kardos et al., 2016). Identification of the causative genetic loci for continuous traits is complex when contrasted to Mendelian loci, as larger sample sizes and precise phenotype scoring methods are required to detect loci (Brien et al., 2022; Kardos et al., 2016; Rockman, 2012). These challenges are highlighted by studies of skin and eye colour variation in human populations, where large sample sizes are required to resolve genome-wide association signals due to small effect sizes and complex ancestries in the datasets (Ju and Mathieson, 2021; Martin et al., 2017; Simcoe et al., 2021).

Here, we use North American *Colias* butterflies to determine the genetic architecture of continuous variation in pterin-based colour in the wild. Pterin pigments are GTP metabolites that are often co-opted to create bright reds, oranges and yellows in a range of phylogenetically diverse animals (Andrade and Carneiro, 2021). As they constitute the red pigments in *Drosophila* eyes, the metabolic pathway has been well-described through mutagenesis screens, although key steps in the pathway, including the hypothesised enzyme that converts xanthopterin to erythropterin, are not yet identified (Vargas-Lowman et al., 2019). Further, there has been limited study of genetic variation in pterin pigmentation in wild populations, with natural allelic variants only identified so far in the mapping of discrete switches among colour morphs of *Podarcis* lizards, and the Alba morph of *Colias* butterflies (Andrade et al., 2019; Woronik et al., 2019).

*Colias* exhibit a range of phenotypes that vary between a closely related species pair. *C. eurytheme* display orange pterins and *C. philodice*, yellow pterins(Morehouse et al., 2007; Wijnen et al., 2007), while females of both species carrying the low frequency *Alba* polymorphism are white due to the suppression of pterin granules in their wing scales (Tunström et al., 2021; Woronik et al., 2019). This species pair is in secondary sympatry in the eastern United States, where they undergo extensive hybridisation in high-density agricultural sites, resulting in continuous variation in their pterin colouration (Gerould, 1943). These visible colour differences are accompanied by a number of other segregating phenotypic differences, including the Mendelian-inherited presence or absence of ultraviolet iridescence (Ficarrotta et al., 2022; Silberglied and Taylor, 1973), as well as variation in size (Grula and Taylor, 1980a), pheromones (Grula et al., 1980; Grula and Taylor, 1979) and mate preference (Grula and Taylor, 1980b). Notably, these phenotypes have all previously been described as sex-linked, indicating that the Z chromosome has a large effect in driving between-species differences. Consistent with a large-X effect (Presgraves, 2018), a recent analysis of between-species differentiation showed a notably large divergence ratio between the Z chromosome and autosomes, though some evidence for diversification on the autosomes was also detected (Ficarrotta et al., 2022).

In this study we examined colour variation in laboratory crosses of *C. philodice* and *C. eurytheme* using an automated image analysis pipeline across 705 butterflies, and found extensive quantitative variation in visible pterin-based colour. Using a previously published 2b-RAD linkage map for 483 individuals from F_2_ and backcross (BC) broods, we mapped two large effect Quantitative Trait Loci (QTL) that interact and jointly explain 70% of the observed wing colour variation, including linkage to an autosome and the Z sex chromosome. We identified two candidate genes that modulate pterin colour output upon CRISPR mosaic knock-out (mKO). Finally, we measured other features of wing morphology including size and pattern elements, finding these traits to similarly be polygenic but with evidence of linkage to the Z chromosome, supporting a model in which the sex chromosomes play an outsized role in the phenotypic diversification between incipient species.

## Results

### High-throughput colour analysis of yellow-orange colour variation in interspecific crosses

We previously generated and genotyped a large cohort of *Colias eurytheme x philodice* laboratory crosses in Amherst, MA in 2000-2002, and used these data to map the genetic basis of ultraviolet colouration, a discrete and male-specific trait (Ficarrotta et al., 2022; Wang and Porter, 2004). This genotype data from 483 individuals in hand, we expanded on this resource to derive wing colour and morphometrics continuous data, by scanning and phenotyping the wings from a total of 705 butterflies, adding additional F_0_ parental populations of *C. eurytheme* and *C. philodice* as well as the offspring from two reciprocal F_1_ crosses, an F_2_ intercross, and two backcrosses of F_1_ males of reciprocal ancestries to *C. eurytheme* females. Automated colour measurements were taken for all four wing surfaces of all individuals. We sampled all pixel values in *L*a*b\** colour space and applied threshold filters to eliminate pixels outside of the yellow-red portion of the spectrum. Noting that the axis with maximum variation was *b\**, we subdivided the *b\** axis into 50 bins (a stable subdivision of the total space), and counted the percentage of pixels which fell within each bin (Weller and Westneat, 2019) (**Figure 1 A-B**). Because of the multidimensional nature of *L*a*b\** colour values, we simply recorded the ‘yellow score’ as the modal bin position (*i.e*. having the most pixels on the colour profile). The yellow scores of the dorsal and ventral forewing, as well as the dorsal hindwing, were highly correlated, while its values showed low variance in the ventral hindwings, where it was decoupled from the three other wing surfaces (**Figure S1A**). As such we used dorsal forewing yellow scores for all further analysis **(Figure 1C-G, Figure 2**).

**Figure 1.**
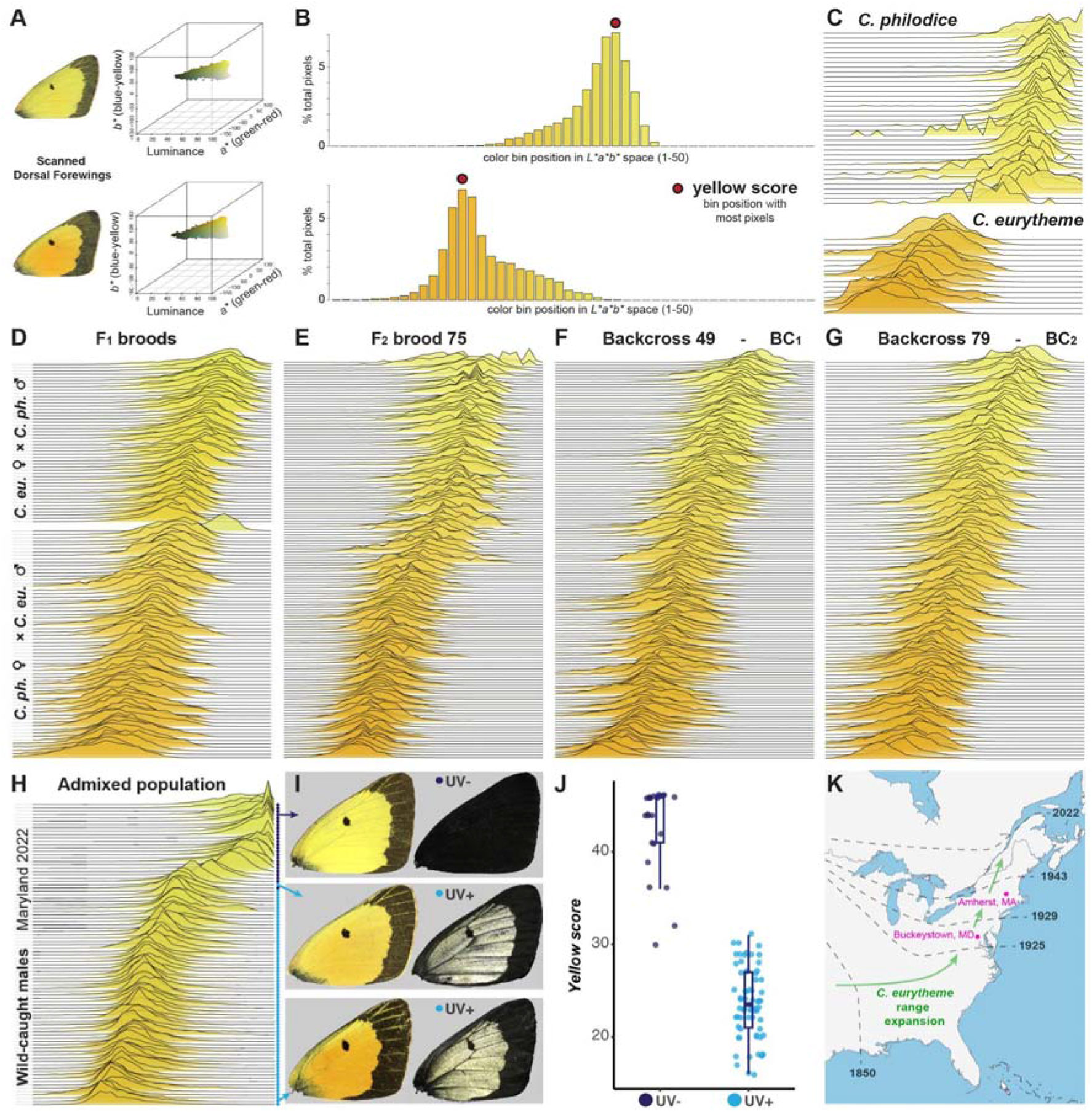
Rapid, repeatable and automated phenotyping of wing colour. (**A**) We used the package *r/colordistance* to extract pixel *L*a*b\** coordinates from scanned wing images, followed by appropriate thresholding to remove background information - here we depict two representative males from the F_2_ brood. We then selected a subset of the remaining pixel range to capture the orange-yellow region of CIELAB colour space, and divided it into 50 bins along the *b\** axis. (**B**) Pixel density histograms for the two individuals shown in A, with colour profile bin positions on the X axis. Bar colours correspond to the average colour of each plotted bin and recapitulate the range of orange-to-yellow pixels of interest in further analyses. The bin with the maximum pixel count is hereafter referred to as ‘yellow score’. (**C-G**) Colour binning profiles of every individual from the offspring of wild-caught individuals (**C**) and indicated broods (**D-G**). (**H-I**) Wild-caught males from an agricultural site where *C. philodice* and *C. eurytheme* fly together and hybridise; the coloured dots indicate the presence or absence of UV iridescence, allowing inference of the Z chromosome genotype (Ficarrotta et al, 2022). (**J**) UV-positive males are consistently oranger while UV-negative males are consistently yellower. (**K**) Summary of the northward expansion of *C. eurytheme* across the Eastern US after 1850, as previously documented (Hovanitz, 1944). The Eastern US comprised part of the native range of *C. philodice* before 1850. Parental admixed populations used for QTL mapping (C-J) originated from Amherst, MA (Summer 2000). Wild-caught males (H-I) were collected in Buckeystown, MD (Aug-Sep 2022).

**Figure 2.**
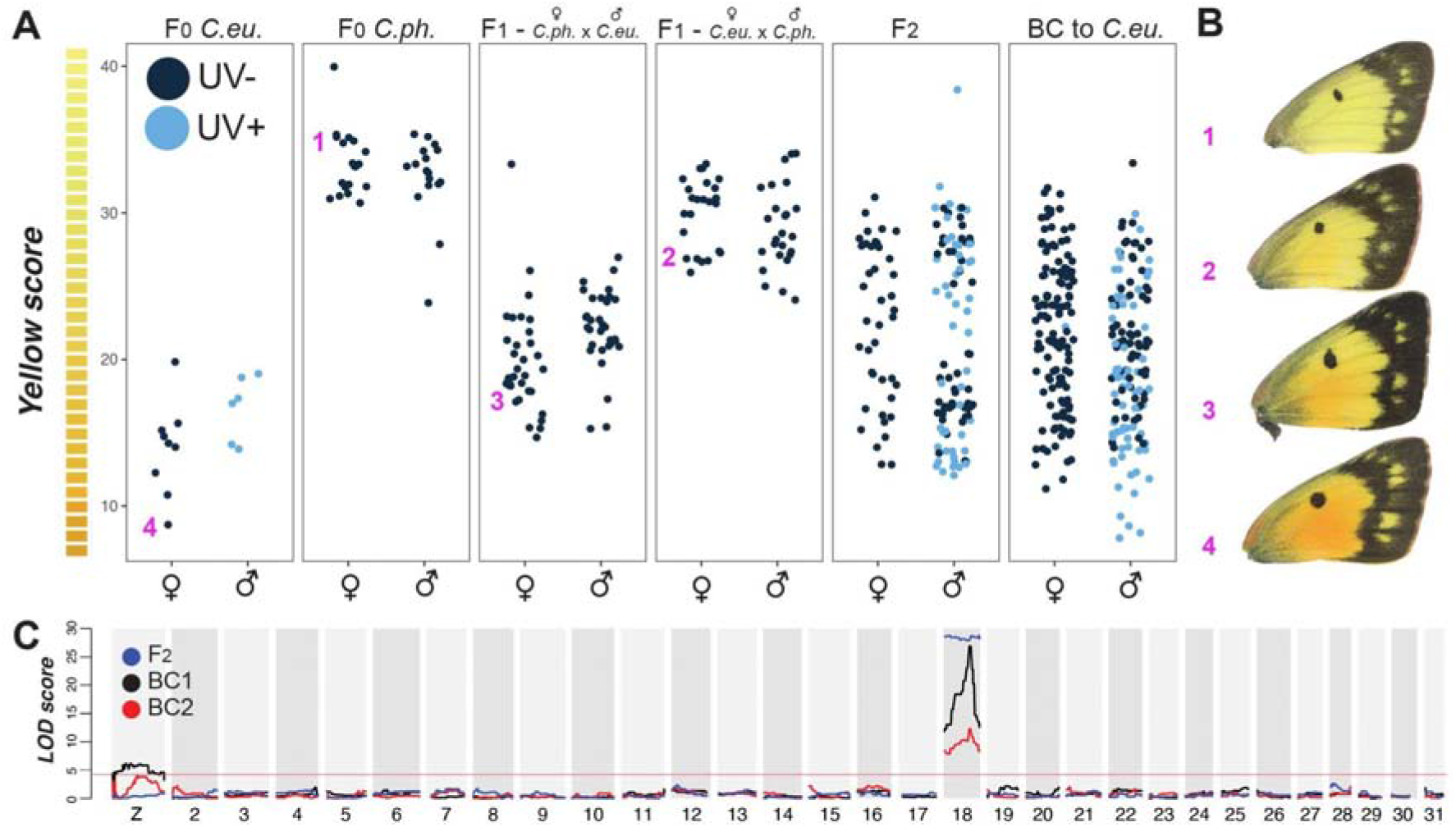
Yellow-orange colour variation is linked to chromosome 18 and the Z sex chromosome. We measured yellow scores for all individuals from one F_2_ and two backcross broods, as well as for a random subset of individuals from grandparental F_0_ and parental F_1_ broods. F_1_ and F_2_ broods show intermediate colour between the parental broods, (**A**). Four exemplar individuals from different parts of the colour range are depicted in (**B**). We used these yellow scores as the input into QTL analyses, and found a LOD interval on chromosome 18 for all three broods, as well as LOD interval on the Z chromosome in both backcross broods (**C**).

As expected, parental F_0_ broods from wild-caught Massachusetts females had colour values at each end of the range (**Figure 1, 2A-B**). Offspring *C. philodice* siblings were yellower (yellow score 32.9±2.5) and *C. eurytheme* siblings were oranger (yellow score 15.1±3.1; Mann-Whitney *U* test, *P* = 2.3 x 10^-8^). The mean colour of F_1_ broods was dependent on the direction of the cross (**Figure 1D**) – offspring with a *C. eurytheme* father were skewed to the orange end of the distribution (skewness of - 0.12), while individuals with a *C. philodice* father were skewed to the yellow end of the distribution (skewness of +0.42). Similarly, F_2_ and BC broods showed an intermediate distribution between the parental phenotypes (**Figure 1E-G**). These trends replicate previous observations and suggest a contribution of the male-inherited Z sex chromosome to colour (Grula and Taylor, 1980a).

Random sampling of genetically unrelated males at a high-density agricultural site in Buckeystown, Maryland exhibited a similar bimodal distribution in yellow score with few intermediates (**Figure 1H-K**). This said, the UV-iridescence characteristic of *C. eurytheme* males is a recessive Z-linked character, allowing us to determine the genotype of the Z chromosome in these individuals from phenotype (Ficarrotta et al., 2022; Silberglied and Taylor, 1973). Wild-caught Maryland UV-positive males (*i.e*. carrying two *C. eurytheme* Z chromosomes) had a yellow score of 23.7±3.8 while wild-caught UV-negative males (either one or two *C. philodice* Z chromosomes) had significantly yellower score of 42.8±4.7 (Mann-Whitney *U* test, *P* = 2.3 x 10^-13^). Taken together, phenotypic data from laboratory crosses and an admixed population in field conditions both suggest the orange-yellow variation is a bimodal species-diagnostic trait, and that its control is at least partially sex-linked. We previously described a large-X effect that caused a strong differentiation of the Z chromosome haplotypes in the Maryland population, limiting genetic resolution by association mapping on this linkage group (Ficarrotta et al., 2022). We thus focused on QTL mapping using laboratory crosses, where both autosomes and the Z chromosome have undergone significant recombination in F_1_ hybrid males (Ficarrotta et al., 2022; Tunström et al., 2021)

### Single QTL scan reveals an autosomal large-effect locus and a sex-linked modifier locus

We calculated an estimate for the effective number of genetic loci required to explain heritability and variance in yellow score using the Castle–Wright estimator, giving an estimate of 3.7 loci; this is likely an underestimate as it assumes all loci have equal effect size, but suggests a model with fewer loci of moderate-to-large effect is likely (Cockerham, 1986; Otto and Jones, 2000). Yellow scores were used as phenotypes for a single QTL analysis, with each brood analysed separately. All three broods had a significant QTL on chromosome 18, with a narrow Bayes-credible interval (**Figure 2C**). Unlike the backcross broods, the F_2_ brood LOD scores on chromosome 18 were flat and the predicted LOD interval incorporated the entire chromosome indicating that no informative recombinants were detected in this brood. The single QTL model also detected a marginally significant, secondary QTL on the Z chromosome in the BC1 and BC2 broods. (**Figure 2D**). It is notable that despite the strong correlation between the Z-linked UV phenotype and yellow score in the Buckeystown wild-caught individuals, the linkage interval with the highest LOD score is autosomal. This, as well as the expectation of additional loci, prompted further dissection of the genetic architecture of colour variation in our mapping panel.

### A two-locus interaction explains 70% of colour variation

To look for interactions and for other, smaller-effect loci, we fit a two-QTL model to each brood separately. For each brood, we identified significant QTL on both chromosome 18 and the Z chromosome, which together explain about 70% of variance in yellow score (**Figure 3A**). At each QTL, individuals with *C. eurytheme* alleles had low yellow scores, individuals with *C. philodice* alleles had high yellow scores, and heterozygotes had intermediate yellow scores (**Figure 3B**). This additive effect is supported by the additive model (plotted below the diagonal in **Figure 3A**), which was significant in BC1 and BC2, but did not reach the genome-wide threshold of significance in the F_2_ brood.

**Figure 3.**
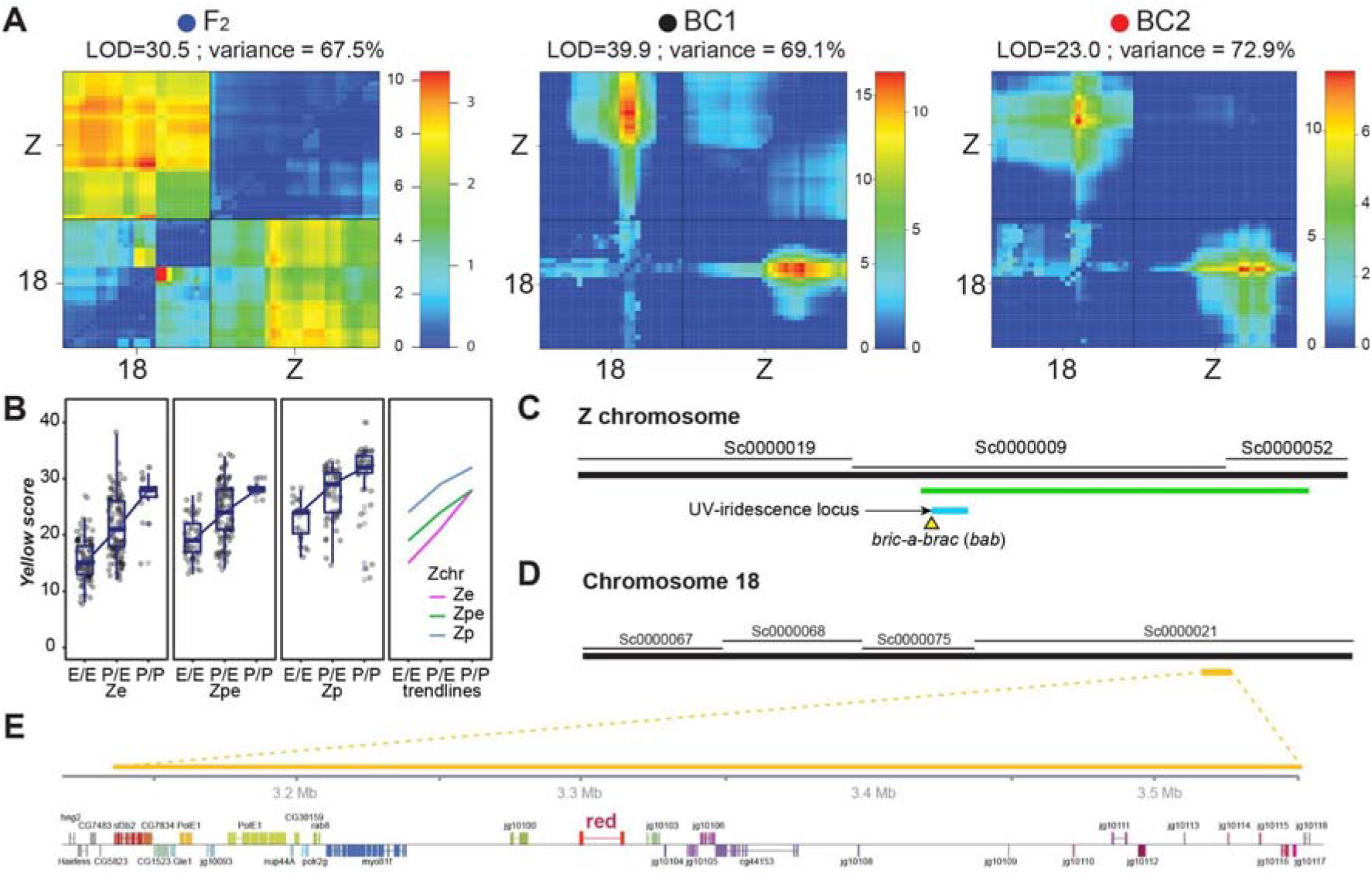
Additive interaction between the LOD intervals on chromosome 18 and the Z chromosome. **A**. Two-QTL models for each brood indicate a significant interaction between the Z chromosome and chromosome 18. For each plot, the top left corner is the full model (two-QTL model plus interaction) and tests for epistasis, whereas the lower right tests for additivity among loci (two-QTL without interaction). Each colour scale indicates two separate LOD scores for epistasis (left) and additivity (right). Significance threshold is set at 4.7 based on previous recommendations (Broman et al., 2003). **B**. Genotype x Phenotype plots showing the relationship between the Z chromosome genotypes (*Ze* and *Zp*: homozygotes or hemizygotes; *Zpe*: heterozygotes) and chromosome 18 LOD interval genotype (x-axis) on yellow score. The rightmost panel overlays the means for the first three panels. **C**. The Z chromosome includes the *U* locus (cyan) locus for UV-iridescence (Ficarrotta et al., 2022). **D-E**. The LOD interval on Chromosome 18 (yellow) is centred on the gene *red Malpighian tubules* (abbr. *red*).

All three broods also show evidence for a significant interaction between the two LOD intervals as shown in the full conditional model (**Figure 3A**, above the diagonal). As indicated by the Genotype x Phenotype plot, when individuals are homozygous for the *C. philodice* allele at the chromosome 18 LOD interval, one copy of the *C. eurytheme* allele at the Z LOD interval makes them more orange, but two copies does not shift this any further. This deviation from the sum of the effects of the two loci indicates an epistatic interaction. To determine if additional smaller effect loci could be detected, we ran multiple-QTL models that accounted for the two detected loci. No additional loci met the genome-wide threshold of significance.

The chromosome 18 LOD interval is 500 kb and contains 29 annotated genes (**Table S1**), including a homolog of the gene *red Malpighian tubules* (*red*). The *red* gene is a compelling candidate gene for pterin pigment variation in butterflies, as mutant and knockdown phenotypes in *Drosophila* and *Oncopeltus* have suggested a role of this gene in the biogenesis of pigment granule formation, including pterinosomes (Francescutti et al., 2022; Grant et al., 2016). No coding variants were detected in whole genome resequenced individuals from Buckeystown, MA.

The Z chromosome LOD interval encompasses a large portion of the chromosome and spans 270 genes (**Table S2**), including the *U* locus interval with the *bric-a-brac* (*bab*) transcription factor gene, which determines male-limited UV iridescence variation in *C. eurytheme* and *C. philodice* (Ficarrotta et al., 2022; Silberglied and Taylor, 1978). Interestingly, we previously reported unusual orange-yellow variation in *C. eurytheme* females following CRISPR mutagenesis of *bab*, prompting a more refined examination of this genetic effect and suggesting that *bab* controls both structural and pigment-based colour differences between the two species.

Lastly, we ran a two-QTL model for the full dataset combining all three broods, and detected a marginally significant interaction between chromosome 18 and chromosome 12 which was not observed in the analyses on separate broods. The chromosome 12 LOD interval incorporates 232 genes and explains 3% of variation in colour (**Figure S2, Table S3**).

### Mosaic knockouts of *red* disrupt pigmentation and intraluminal scale organisation

In order to test a role of *red* in wing pterin modulation, we used CRISPR mutagenesis targeting a coding exon of *red* to generate null, mosaic mutant clones in G_0_ injected individuals. A total of 305 syncytial embryos were collected from wild-caught *C. eurytheme* females and injected within 4 h after egg laying (AEL). From these injections, we obtained 27 emerged adult butterflies, 20 of which (74%) showed visible colour changes in mutant clones (**Figure 4, Table S4**). Both ventral and dorsal wing surfaces were affected, and mutant clones showed a gain of broad-spectrum (non-iridescent) ultraviolet reflectance (**Figures S3-4)**, suggesting a decrease of UV-absorbing pterin pigments in mutant scales (Wijnen et al., 2007; Wilts et al., 2017). Dorsal hindwing discal spots, which normally display a deep orange colour regardless of the species and sex, showed the most marked colour shifts among crispants (**Figure S5**). Elsewhere on the wing, and most markedly on dorsal surfaces, the pale colour of mutant clones somewhat resembled Alba female morphs but with a higher chromaticity (**Figure S4A-K**), implying that *red* KOs do not fully suppress pterin granule formation as observed in *Alba* genotypes (Woronik et al., 2019). To better characterise the effects of *red* loss-of-function on wing chromaticity, we digitised crispant wings in constant conditions and then analysed wild type and mutant clones in immediately adjacent patches of wing in CIELAB colour space (**Figure 4I-K**). Wildtype populations show little variation in luminance (*L\**), but *C. eurytheme* has higher *a\** values (*i.e*. they are redder) and *b\** values (*i.e*. they are also yellower) than *C. philodice*. Crispant clones consistently showed a small reduction in *a\** and remained above the *C. philodice* states while *b\** values fell under the *C. philodice* values. This overall reduction in chromaticity suggests a decrease in the pterin content of *red* crispant wings, akin to but less drastic than in Alba females (Woronik et al., 2019).

**Figure 4.**
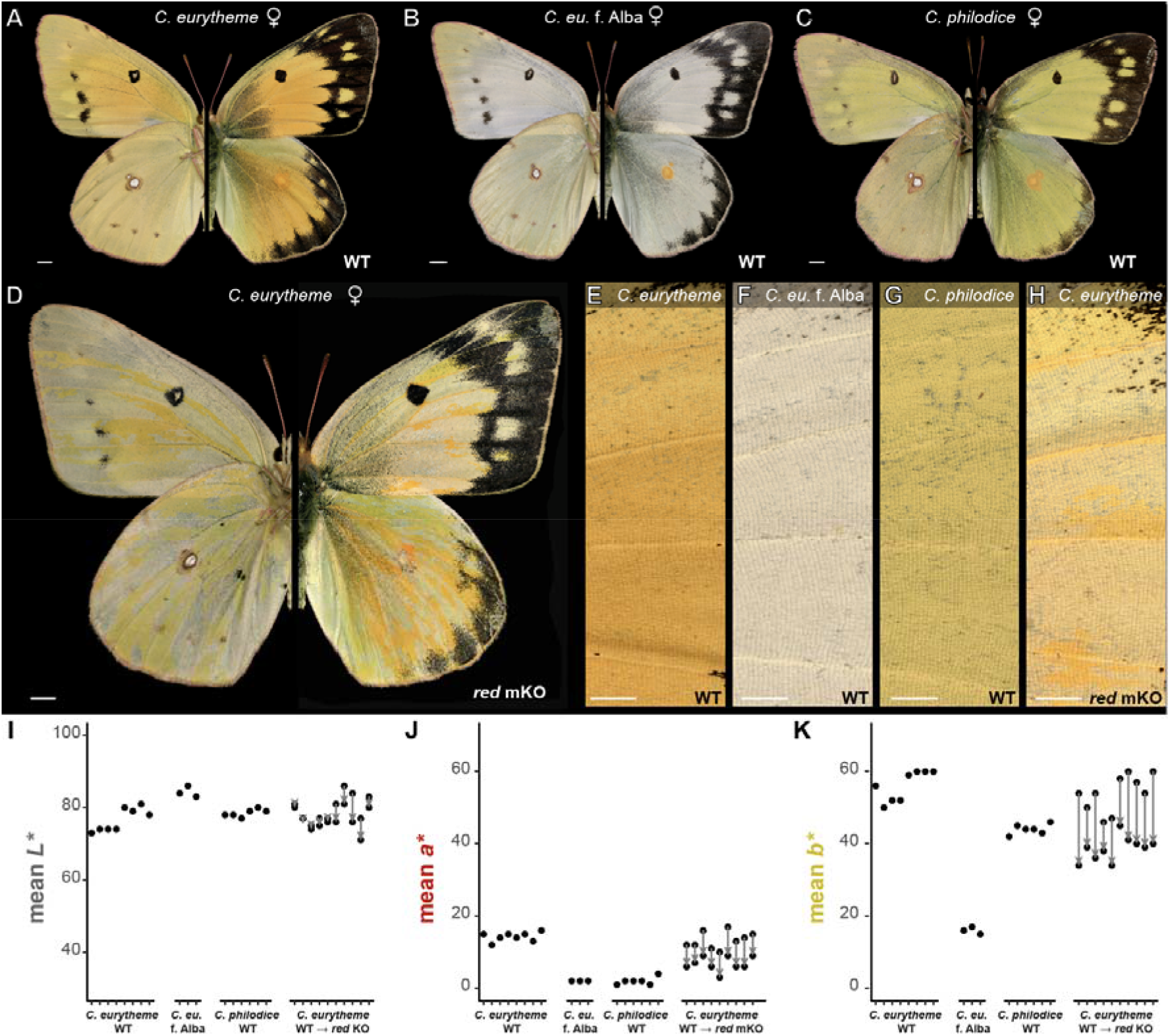
Wing colouration effects of *red Malpighian tubules* CRISPR mKOs. **A-C.** Ventral (left) and dorsal of wild-type (WT) females representative of admixed *C. eurytheme* and *C. philodice* populations, here sampled from Maryland. **D.** *C. eurytheme* G_0_ mosaic crispant for the *red* gene with discoloured streaks (mutant clones), imaged in identical conditions to the wild-type controls. **E-H**. Side-by-side comparisons of dorsal forewing regions show reduced orange colouration in *red* KO clones (panel H) relative to *C. eurytheme* WT clones; compare to Alba individuals, which completely lack pterin granules. **IK**. Comparison of mean *L\** (luminance), *a\**(green to red), and *b\** (blue to yellow) pixel values sampled from dorsal WT forewings, and from adjacent WT vs. *red*-deficient clones from 10 *C. eurytheme* crispants, contrasted to Alba individuals which lack pterins. Scale bars: A-D = 2,000 μm; E-H = 1,000 μm.

In pierid butterflies, pterin pigment granules (pterinosomes) can be directly visualised by scanning electron microscopy. Pterinosomes normally form oblong ellipsoid structures attached along the transversal trabeculae of the upper scale surface, and suspended in the hollow scale matrix (Morehouse et al., 2007). This organisation was disrupted in *red* crispant scales, which showed a normal organisation of the scale outer structures, but a chaotic mesh of interwoven structures in their inner part (**Figure 5**). Pterinosomes were apparent but scarce and disorganised, likely explaining the residual chromaticity in these scales, and fused with other chitinous elements. Under the hypothesis that pterinosomes are lysosome-related organelles that derive from the scale endomembrane system (Figon et al., 2021b; Ghiradella, 2010), these phenotypes are consistent with a disruption of intraluminal organelle maturation or trafficking.

**Figure 5.**
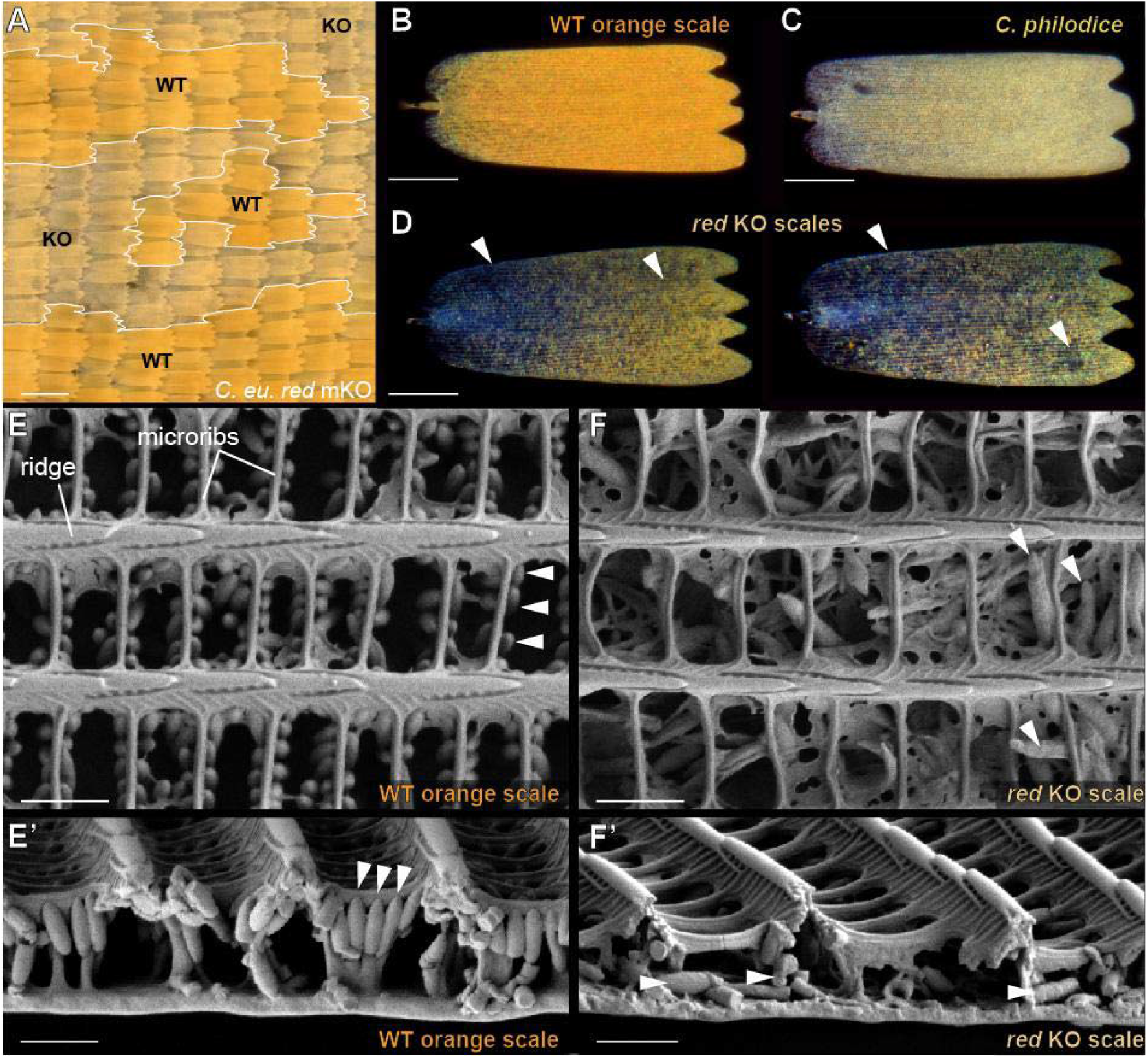
Disorganisation of internal scale structures in *red* crispant scales. **A.** Scale-level colouration of *red* mKO clones relative to wild-type (WT) phenotypes in *C. eurytheme*, here in dorsal forewing regions. **B-D**. Reflected polarised microscopy of individual dorsal forewing scales from a *C. eurytheme* female (**B**), a *C. philodice* female (**C**), and a female *C. eurytheme red* crispant clone (**D**). Scales from panels B and D are from adjacent clones in the same crispant individual. Scales deficient for *red* have poor refringence and show an overall lack of pigment density (arrowheads). **E-E’**. SEM top (**E)** and cross-sectional (**E’**) views of WT orange scales. Pterin granules (arrowheads) are visible as oblong structures attached to the microribs, the structures that form transversal bridges between the ridges of the scale upper lamina. **F-F’**. Defects of the inner scale lumen in *red* crispant scales, amidst a normal upper surface. Pterin granules (arrowheads) are improperly formed, fail to attach to microribs, and are randomly arranged in a disorganised scale matrix. Scale bars: A = 100 μm; B-D = 50 μm; E-F’ = 1 μm.

### The UV-iridescence switch gene *bab* is also a candidate gene for pterin variation

Our two-QTL scan identified a large portion of the Z chromosome as a secondary locus interacting with chromosome 18, with a confidence interval that includes the U locus, previously described in Ficarrotta et al. (2022), which contains the causative gene *bab*. *C. eurytheme* and *C. philodice bab* crispant clones display spectacular gains of UV-iridescence across all wing surfaces and in both sexes. Interestingly, we previously noticed that while *bab* mKOs do not yield effects on human-visible colour in males, *C. eurytheme* crispant females displayed noticeable heterogeneity in visible colour (Ficarrotta et al., 2022). We re-examined this phenomenon and found the *bab* loss-of-function clones resulted in context-dependent effects on several colour traits in *C. eurytheme* females (**Figure 6**).

**Figure 6.**
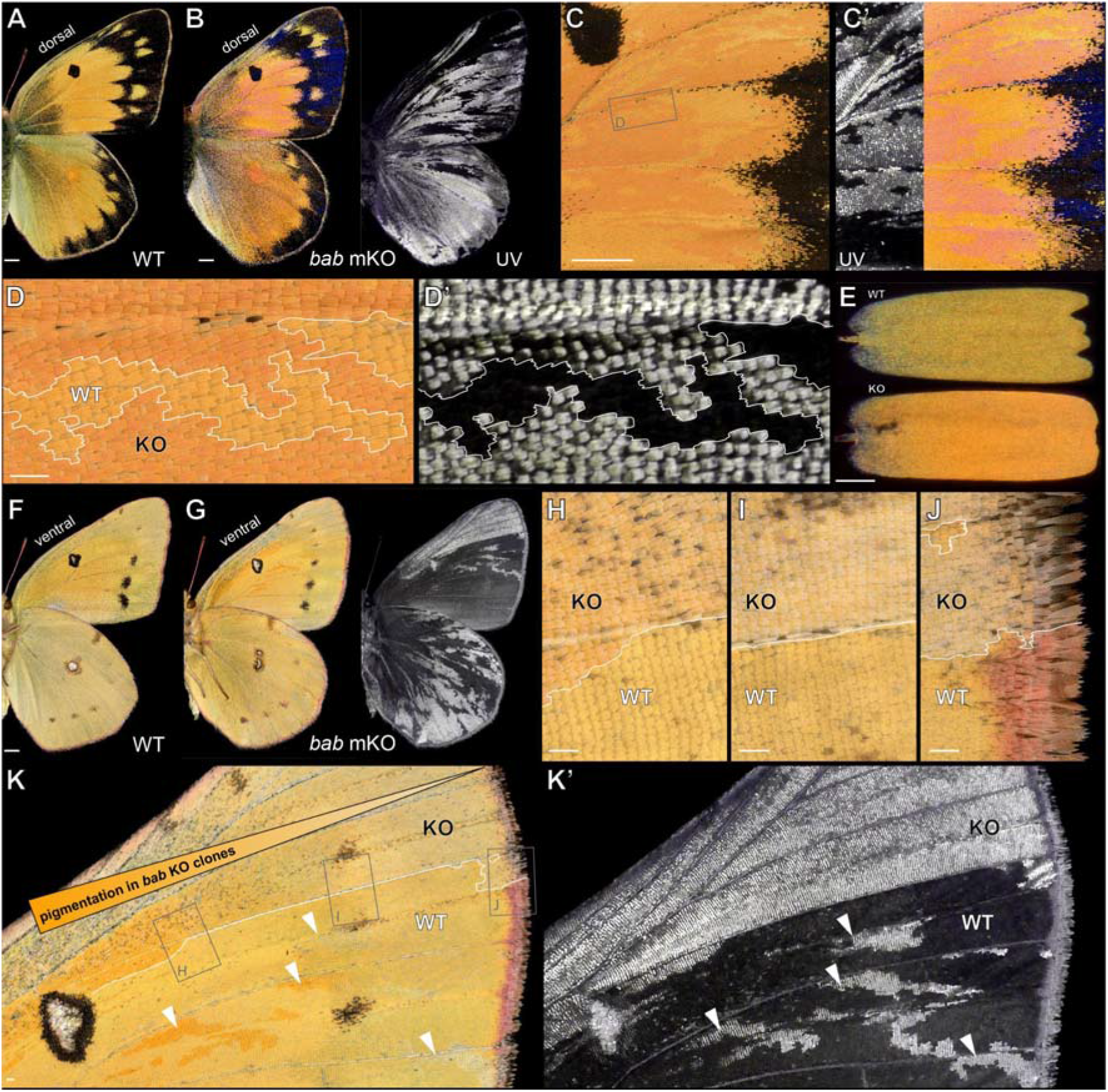
CRISPR mutagenesis of *bab* can affect both UV-iridescence and pterin pigmentation. **A.** WT *C. eurytheme* female, dorsal view. **B-C’**. Dorsal views of a *bab* crispant *C. eurytheme* female (Ficarrotta et al., 2022). Crispant clones lacking *bab* display ectopic UV-iridescence in all regions, ectopic blue iridescence over melanic regions, and colour shifts in orange regions, with the two latter effects only observed in crispant females. C shows the visible-spectrum colouration with normal light incidence (0°), while C’ reveals the structural iridescence effects in the UV-A and visible ranges observed with a 30° light incidence. **D-D’.** Magnified views of WT and KO *bab* crispant clones with 0° incidence. **E**. Polarised reflective microscopy of adjacent scales from across a *bab* crispant boundary. **F**. WT *C. eurytheme* female, ventral view. **G-K’**. Ventral views of a *bab* crispant *C. eurytheme* female. On the ventral surface, the effect of *bab* loss-of-function over pterin pigmentation changes over the proximo-distal axis. Scale bars: A-B, F-G = 2,000 μm; C-D, H-K = 200 μm; E = 50 μm.

Observation of wings with a 30° light incidence highlighted a gain of an iridescent colour, visible as a blue sheen on melanic scales and as a violet-pink sheen on orange scales (**Figure 6C’**). This effect is limited to females, and is likely an effect of UV-signal overflow in the visible spectrum following the gain of densely stacked ridges on the surface of the UV-iridescent scale type. As this iridescent colour is angle dependent, we imaged these wings at a 0° incidence as well as with polarised light reflected microscopy, to rule out this ectopic iridescence as having an effect on the appearance of pterin-based colour (**Figure 6C**). Because mutant *vs*. WT clones maintained marked differences in orange-yellow pigmentation with these observation techniques, we infer that *bab* KOs results in the modulation of pterin content in females, separately from its role in structural colour.

Of note, *bab* crispant clones are marked by ectopic UV-iridescence, allowing us to precisely delineate the boundaries of WT vs mutant wing scales. On the dorsal wings, we found that *bab*-deficient clones show an increase in orange pigment compared to adjacent WT clones (**Figure 6D-E**). The analysis of ventral surfaces revealed a more nuanced picture, because the specific effect of *bab* mutagenesis on red pigmentation varied across the proximo-distal axis (**Figure 6F-K’**). In proximal regions, *bab* mutant clones showed redder pigmentation relative to adjacent WT clones, while more distally, they were comparatively lighter, including in the most distal section of the wing where marginal scales that normally display pink pterins (Fenner et al., 2022) were depigmented. In addition, *bab* mKOs resulted in an elongation of silver scales and darker melanisation in the ventral discal spot (**Figure S6**).

In summary, CRISPR knock-outs revealed *bab* is required for normal pterin pigmentation in females in a complex fashion, with *bab-*deficient clones showing increased orange pigmentation on dorsal surfaces, and varying effects across the proximo-distal axis on ventral sides. Allelic variation of the *bab* locus may thus explain all of the colour variation linked to the Z chromosome LOD interval, possibly through the direct activation or repression of pigment pathway components – similarly to its role in shaping *Drosophila* abdominal pigmentation via other interacting QTL (Castro et al., 2018; Kopp et al., 2003; Rogers et al., 2013). Alternatively, the sex-linked secondary QTL may be explained by other genes independent of *bab*. Traditional QTL mapping may be limited to test this possibility, as it might require the generation of many large recombinant broods controlling for *red* allelic states to properly extract a narrow signal on the Z chromosome.

### Wing size is heritable and sex-linked

Previous work showed that *C. eurytheme* were consistently larger than *C. philodice*, and that this size difference was heritable and sex-linked (Grula and Taylor, 1980a; Porter and Levin, 2007). We quantified wing size, along with a number of other morphometric measurements including features of the discal spots and the marginal bands, from all 705 individuals (**Figure S7**). Principal component analysis of all measures showed separation between UV-positive and UV-negative males in the first component, once again indicating a likely contribution from the sex chromosomes. Of note, we replicated a small but significant difference in wing length between females of the two parental species (p = 0.03, Wilcoxon test) (**Figure 7**).

**Figure 7.**
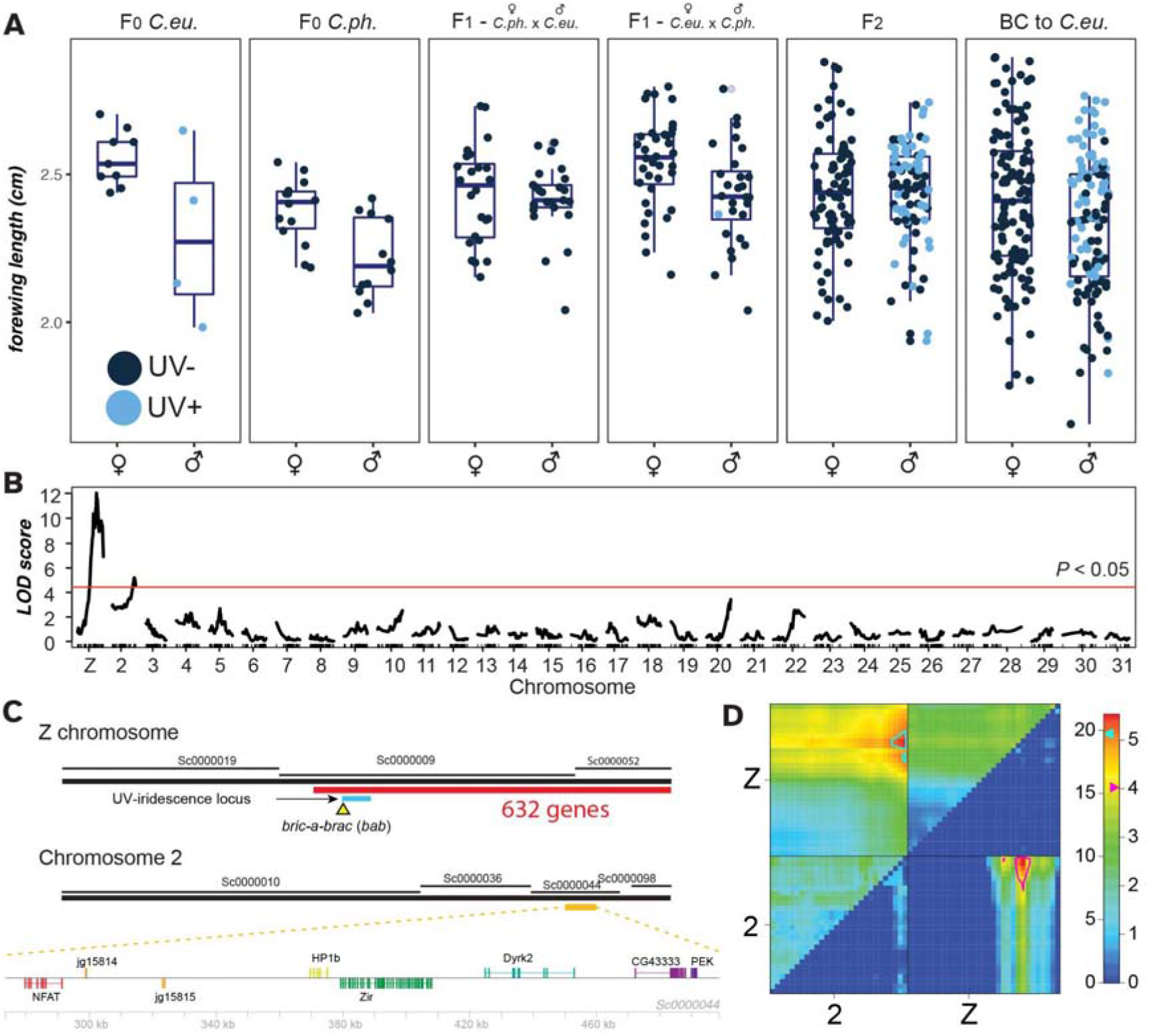
Two LOD intervals associated with wing size. **A.** Variation in wing length in crosses. **B-C.** A single-QTL model detects two significant loci (B), which incorporate large portions of chromosome 2 and the Z chromosome (C). **D.** Support for an additive two-QTL model that explains 18% of the variation in size, with LOD scores for the full model on the upper left and for the additive model on the lower right.

The morphometric features we measured could have been controlled by the same or linked loci and might not be independent variables, so in order to determine the co-inheritance of these traits we performed a hierarchical clustering analysis to look for covariance between measurements. Our measured traits optimally split into 5 clusters, one each corresponding to size; forewing discal spot; hindwing discal spot; forewing and hindwing black margin thickness; and wing aspect. We then used the latent variable for each of these trait clusters for QTL analysis (**Figure S8-9**). Two significant LOD intervals for size were detected, on the Z chromosome and on chromosome 2 (**Figure 7**). In a two-QTL model, an additive interaction between the two loci explained 18% of variation in size. Additional significant QTL were also detected for the hindwing discal spot, the forewing discal spot, and the thickness of the black margin (**Figure S8**).

The Z-linked LOD interval for size incorporates around half of the chromosome and includes 638 genes, including the entirety of the LOD interval for colour described above (**Table S2**). The LOD interval for size on chromosome 2 was more compact, including 8 genes (**Figure 7C**, **Table S5**). Further linkage analysis or association studies would be required to allow the determination of candidate loci for variation between *C. eurytheme* and *C. philodice*; this trait is likely to be highly polygenic, reflecting their divergent life-history strategies for size and rate of growth (Grula and Taylor, 1980a; Porter and Levin, 2007).

## Discussion

### Phenotyping continuous colour variation

We were able to rapidly and objectively score continuous colour in many individuals, permitting us to identify the underlying genetic architecture of an interspecific variable trait. There are significant challenges for these types of quantitative analyses that make this approach necessary; the human eye may not always detect all meaningful variation. Objective measurement of colour may reveal real, mappable signals that are indistinguishable in the human vision model; for example, recent analysis of tiger moth wings has shown that automated measurements can detect heterozygotes where humans cannot, highlighting the importance of objective measures (Nokelainen et al., 2022). This is compounded by the requirement for large numbers of individuals for the mapping of quantitative traits; as we continue to map continuous traits in future, we will begin to encounter data sets with many thousands of individuals; to cope with big-data phenomics, we need an increased focus on developing methods that can allow rapid phenotypic characterisation. Approaches like the one used here for colour measurement should be scalable in order to deal with very large and complex data sets.

### The genetic architecture of yellow-orange colour variation is oligogenic

We used interspecific laboratory crosses of *C. eurytheme* x *philodice*, two sulphur butterfly species that occur in secondary sympatry, to shed light on the genetic architecture of continuous wing traits, and used a high-throughput colour quantitation method to map the loci underlying the yellow-orange colour characteristic of these species. We found that two large-effect loci and one small effect locus, together explain 73% of the heritable variance in yellow score and indicate a mostly oligogenic architecture. This result contrasts with general expectations that many loci of smaller effect should underlie quantitative trait variation (Boyle et al., 2017; Rockman, 2012) – as illustrated (using similar methodology and traits) by three additive QTL that explained only 8% of the heritable variation in ornamental orange colouration in stickleback fishes (Yong et al., 2016). Our epistatic QTL detected in the two-QTL scan were highly significant, though we may overestimate due to the Beavis effect (Xu, 2003). This means that 27% or more of the colour variance remains unaccounted for in our mapping panel, and it would require high statistical power from large crosses to identify further underlying small-effect loci (Mackay et al., 2009; Rockman, 2012; San-Jose and Roulin, 2017).

Finally, yellow-orange variation is known to be seasonal (Gerould, 1943; Hoffmann, 1974; Remington, 1954) and correlated to a melanic trait that is influenced by day length across many pierids (Jacobs and Watt, 1994; Kingsolver, 1995; Kingsolver and Wiernasz, 1991). Here, our mapping approach controlled for Genotype x Environment effects as all broods were reared under constant conditions, but we expect variation in nature to include a more complex architecture, including small-effect loci that may modulate the responsiveness of chroma to photoperiod, temperature, or larval nutrition (Beldade et al., 2011; Lafuente and Beldade, 2019).

### *red Malpighian tubules* – a candidate regulator of pterinosome maturation

In this study, mapping and mosaic knock-out experiments implicated the homolog of the fly *red* gene as a large effect QTL controlling colour variation between sympatric *Colias* species. This gene encodes protein with a conserved LysM domain, although its molecular function has seldom been studied. The *Drosophila* mutant strain *red Malpighian tubules* (*red*) was described in 1954 (Oster, 1954) and has been used as a marker gene (Henikoff, 1979; Breen and Harte, 1991), but its molecular annotation to the *red* coding gene was only recently described (Grant et al., 2016).

This said, several lines of evidence posit *red* as a gene involved in the trafficking and maturation of pigment granules, which are endosomal compartments known as Lysosome-related Organelles or LROs, and are universally involved in pigment processing throughout animals (Figon et al., 2021b, 2021a; Futahashi and Osanai-Futahashi, 2021). The *D. melanogaster red* mutant shows an accumulation of red ommochromes in the Malpighian Tubules (MTs), and a reduction of ommochromes and pterins in the eyes (Aslaksen and Hadorn, 1957; Ferré et al., 1986; Wessing and Bonse, 1966). In the milkweed bug *Oncopeltus fasciatus*, RNAi-mediated knockdown of *red* causes a reduction in pterins in pigmented cuticles, legs, and abdomens, as well as a reduction of ommochromes in the eye (Francescutti et al., 2022). Wing pigmentation is primarily pterin-based in *Colias*, with no contribution from ommochromes (Wijnen et al., 2007), but the dual effects on both ommochromes and pterins in *Drosophila* and *Oncopeltus* suggest that *red* acts at the level of a biological process that is shared between these pathways. Importantly, *red* mutant MTs show aberrant vesicular phenotypes that imply a failure of pigment granules to properly mature and excrete their content in the lumen (Wessing and Bonse, 1966). These effects are reminiscent of other eye colour mutant genes of the ‘granule group’, that are required for the biogenesis of LROs, including ommochromasomes and pterinosomes (Figon and Casas, 2019; Futahashi and Osanai-Futahashi, 2021; Shoup, 1966). In our SEM analysis of *red* crispant scales, we observed defective and disorganised pterinosomes amidst otherwise normal scale morphologies (**Figure 5**). While further work is needed to establish the cellular roles of this gene, these data collectively suggests that *red* is involved in the trafficking or maturation of pigment granules, including pterinosomes and ommochromasomes.

In addition, a few observations lead us to speculate that *red* may regulate pigment granule activity by interacting with the vacuolar pH of pigment granules. First, the unique pigment profile of *red* mutant flies is strikingly similar to the ones observed with two eye-colour mutant alleles of the *chocolate (cho) / VhaAC39* gene (Ferré et al., 1986; Grant et al., 2016; Tearle, 1991), which encodes a v-ATPase proton pump essential for endosomal acidification and membrane trafficking in LROs (Allan et al., 2005; Sun-Wada et al., 2003; Yan et al., 2009). Second, homologs of Red contain a LysM domain, and it is their only annotated feature (Francescutti et al., 2022; Grant et al., 2016). The molecular function of this domain is poorly understood in insects, but there is mounting evidence in vertebrates that the LysM domains of several genes interact with the v-ATPase to modulate vacuolar pH across a variety of endosomal organelles (Merkulova et al., 2015; Castroflorio et al., 2021; Eaton et al., 2021; Tan et al., 2022). There is precedent for a role of pterinosome acidification in modulating the red vs. yellow states of squamate pigment cells (Saenko et al., 2013), and the roles of melanosomal pH and v-ATPase activity in fine-tuning melanin content are well established in vertebrates (Ramos-Balderas et al., 2013; Wakamatsu et al., 2021). Thus, while future work will be needed to dissect the roles of insect *red* and *cho* in v-ATPase function in pigment granules, we speculate that *red* regulates pterin composition by modulating the pterinosome vacuolar pH in *C. eurytheme* and *C. philodice*.

### Evidence for a ‘species-defining’ sex chromosome

The North American sulphur butterflies *C. philodice* and *C. eurytheme*, co-exist in sympatry while maintaining effective reproductive barriers (Wang and Porter, 2004), and the Z sex chromosome plays a disproportionate role in keeping these species distinct, an effect known as the Large-X effect (Presgraves, 2018). Moreover, the Z chromosome has a remarkably high level of divergence compared to autosomes (Ficarrotta et al., 2022), and it also acts as a large linkage group coupling key reproductive barrier traits such as asymmetric hybrid female sterility, both male pheromone signal and female preference, as well as male UV signal and female preference; it thereby acts as a cluster of loci involved in incipient speciation (Grula et al., 1980; Grula and Taylor, 1980b, 1980a, 1979). The UV colouration signal, a recessive marker that distinguishes *C. eurytheme* males from hybrids and allows conspecific females to find compatible gametes, is explained by allelic variation of the *bab* transcription factor, a suppressor of UV scales that is specifically de-repressed in a small fraction of cells in iridescent males.

Here, we showed that two additional traits that vary between this species pair, pterin colouration and wing size, are also linked to the Z chromosome. CRISPR knock-outs implicate a role for *bab* in orange-yellow pigmentation. It thus represents a good candidate gene for regulating interspecific variation, though given the large size and gene content of the mapped QTL, the causative variants may in fact be associated with another gene in the interval. A sexual signalling role of dorsal visible colour (orange vs. yellow) is unlikely in males, and untested in females (Silberglied and Taylor, 1978). While this is also untested, wing chroma differences may have been shaped by ecological adaptation to climatic differences in the native ranges of *C. eurytheme* vs *C. philodice* (Hovanitz, 1944), as suggested by the seasonal plasticity of this trait (Fenner et al., 2022; Gerould, 1943). Wing size differences (a proxy for total body size) are likely to reflect the divergent life history of these butterflies, with *C. philodice* reaching pupation faster than *C. eurytheme* in spite of identical growth rates on their introduced host plants (Porter and Levin, 2007).

The accumulation of LOD intervals for species-defining traits on the Z chromosome suggests that the Z chromosome could be behaving like a chromosome-wide supergene. It is possible that few linked large-effect loci under strong selection, plus the 0.75x recombination rate on sex chromosomes vs. autosomes, can more easily overcome the effects of introgression in comparison to autosomes (Fraïsse and Sachdeva, 2021; Wilson Sayres, 2018), as corroborated by the strong signature of Z-chromosome differentiation between the two *Colias* species in sympatry. On the other hand, we have described additional autosomal loci that have persisted in divergence between the two species, including for size, colour (*i.e. red*), and additional phenotypes (**Figure S8**). This coupling between the Z chromosome and autosomes could represent the remnant of eroding barrier loci, or be the a signal of the persistence of stable ecomorphs maintained by sexual selection (Butlin and Smadja, 2018; Unbehend et al., 2021). The resulting genetic architecture, with comparatively few loci contributing a large effect to a continuous trait, bears similarity to other described cases of colour variation, including blue iridescence in butterflies and plumage colour in capuchino seedeaters where a few large-effect loci explain a high fraction of variance (Brien et al., 2022; Estalles et al., 2022), while standing in contrast to other cases where many smaller effect loci predominate, including the white head patch of *Ficedula* flycatchers (Kardos et al., 2016). This adds to a growing picture of the genetic control of continuous traits.

## Acknowledgements

We thank Baiqing Wang for previously generating mapping broods (Wang, 2005), Vincent Ficarrotta for previously generating *bab* crispants used in this study (Ficarrotta et al., 2022); Scott Barao and Hedgeapple Farm (Buckeystown, MD) for providing access to alfalfa fields; the GW Nanofabrication and Imaging Center for SEM access; Rachel Canalicchio and the Wilbur V. Harlan Greenhouse staff for host plants; and the Research Technology Services at the George Washington University for operating the High Performance Computing Cluster (MacLachlan et al., 2020). We thank the Martin lab for their comments on this paper.

This work was funded by the National Science Foundation (IOS-1755329), and Harlan Foundation summer fellowships to OBWHC and LSL.

## Author contributions

Conceptualization, JJH, AHP and AM; Investigation and Formal Analysis, JJH, CMF, OBWHC and LSL; Data Curation, OBWHC, MAN and DJL; Visualization, JJH and AM; Writing, JJH, AHP and AM.

## STAR Methods

### KEY RESOURCES TABLE

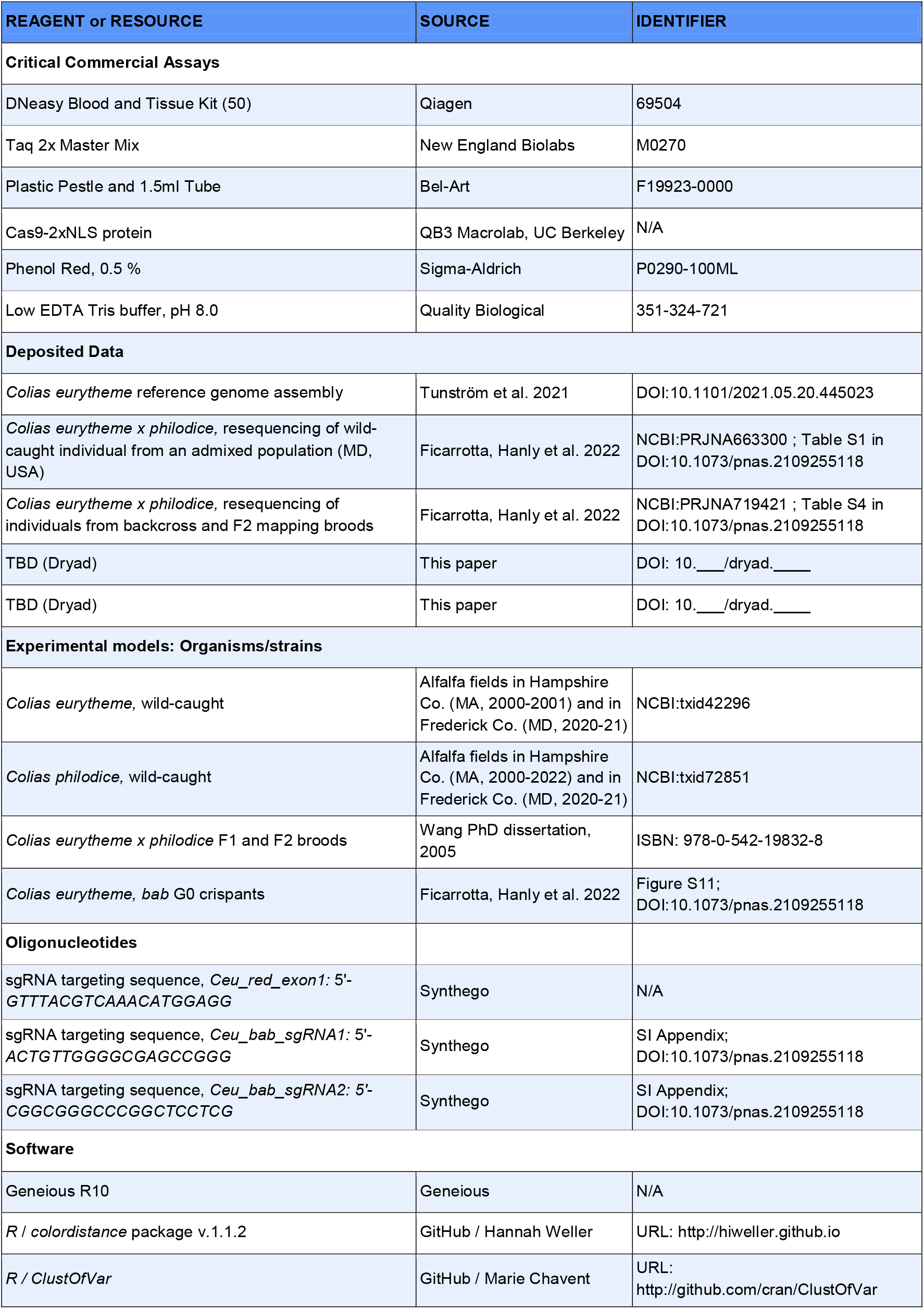
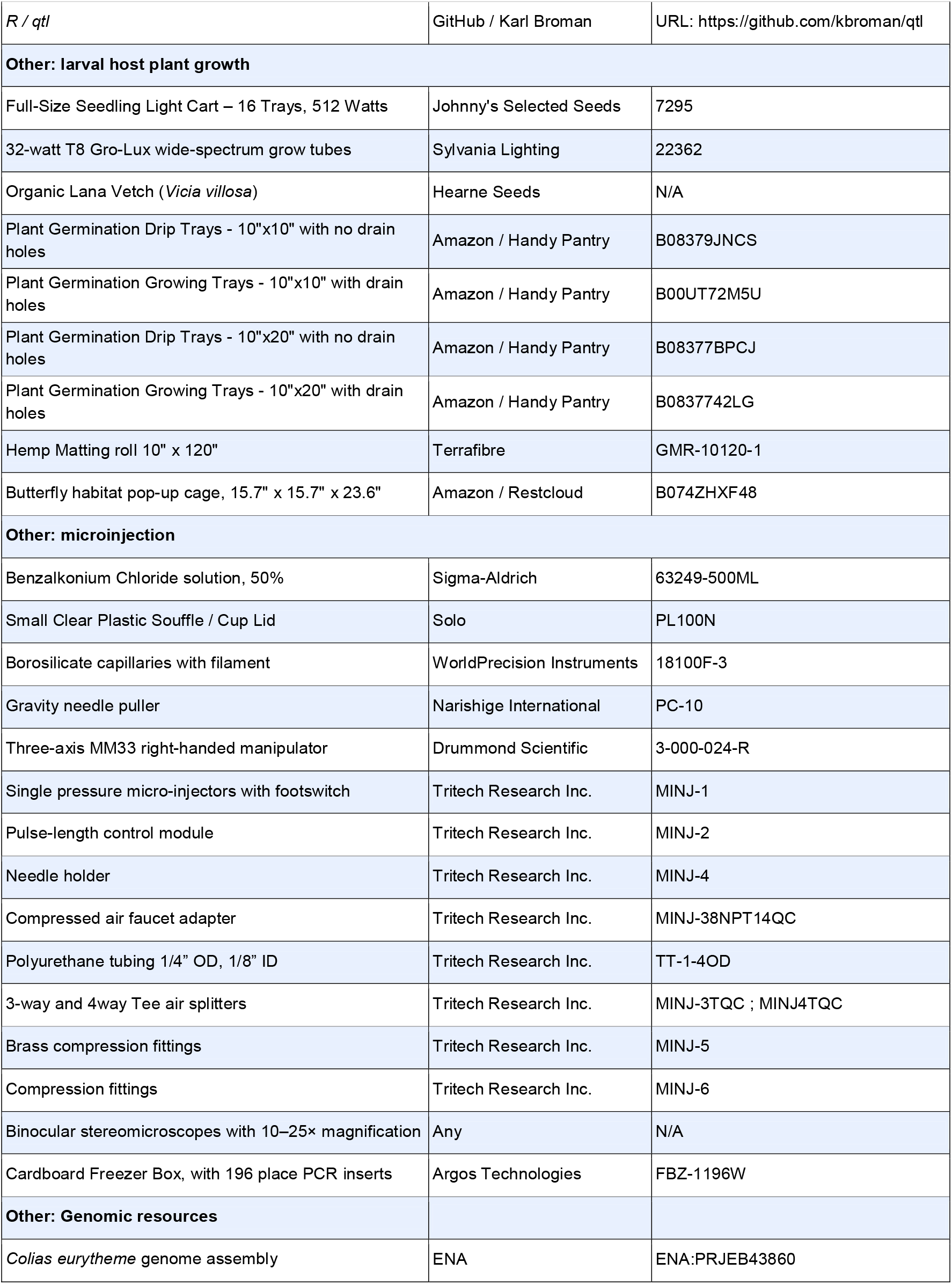

### RESOURCE AVAILABILITY

#### Lead contact

Further information and requests for resources, protocols and reagents should be directed to and will be fulfilled by lead contact, Joseph Hanly

#### Material availability

Wings and DNA samples are stored in the laboratory of Arnaud Martin, and accessible for loan upon request, within the US.

#### Data and code availability

- Experimental model organisms/strains are listed in the key resources table.
- Code and images used in colour analyses have been deposited at Dryad and are publicly available as of the date of publication. DOIs are listed in the key resources table.
- Other data needed for re-analysis will be shared by the lead contact upon request.

### EXPERIMENTAL MODEL AND SUBJECT DETAILS

#### Butterfly mapping broods

Interspecies F_1_, F_2_ and backcrosses between *C. eurytheme* and *C. philodice* were generated in 2000-2001 from wild individuals caught in and around Amherst, MA, as previously described (Wang, 2005; Wang and Porter, 2004). A former publication describes DNA extraction procedures and 2b-RADseq linkage maps for one F_2_ brood and two backcross broods (Ficarrotta et al., 2022; Tunström et al., 2021). All broods were reared under summer conditions (27 °C and LD 14:10 h) on fresh alfalfa cuttings placed in large Petri dishes in two growth chambers. Dishes were moved within and among growth chambers at 3-day intervals to randomise environmental growth chamber effects. Adults were stored in labelled glassine envelopes at −80 °C shortly after emergence until thorax tissue sampling for DNA extraction in 2018.

#### Butterfly mosaic KOs

For CRISPR mutagenesis, *C. eurytheme* gravid females were wild-caught on organic alfalfa fields in Buckeystown, MD (kind permission of Dr. Scott Barrao, Hedgeapple Farms) before being brought in the lab for oviposition on alfalfa cuttings (Ficarrotta et al. 2022). Larvae were reared on hydroponic trays of 7-21 d old vetch sprouts contained in large collapsible cages, in a temperature controlled greenhouse maintained at 27-29 °C and with an automated misting system, in August-September 2020 (*bab* KOs) and June-September 2021 (*red* KOs). Total developmental time at 27-29°C is about 26 d, with an average egg hatching time of 70 h AEL, and an adult emergence time of 132 h after pupa formation. Emerged adults were pinned or their wings stored in glassine envelopes for further imaging.

#### Larval host plants

Alfalfa plants (*Medicago sativa*) were grown as mature plants in pots containing a soil mix with vermiculite. Fresh plant cuttings with young leaves were used for larval rearing, or placed in a water cup for oviposition. Dry seeds of Lana hairy vetch (*Vicia villosa* var. Lana) were weighed, soaked overnight in water for rehydration, spread on water-saturated hemp fibre mats placed in hydroponic trays at a density of 7.5 g / in^2^ (dry weight), germinated in the dark for 72 h with occasional misting with water, fertilised with a 200 ppm N solution (1 g / L of NPK 20:20:20 fertiliser), and sprouted on a seedling light cart equipped with 32 W T8 wide-spectrum tubes for 72 h before removal from the grow lights. Hydroponic trays were occasionally bottom-watered with a 50 ppm N solution and provided to larvae for feeding within 14 d after germination.

### METHOD DETAILS

#### Colour imaging

For QTL mapping, we scanned the dorsal and ventral surface of forewings and hindwings from the three previously described broods, plus wings from four additional broods, with an Epson Perfection V600 scanner with a 24-colour correction card and greyscale standards.

For colour quantification in *red* CRISPR experiments, dorsal wing regions of interest situated between the M_1_-Cu_2_ veins were digitised with a Keyence VHX-5000 microscope, and VH-Z100T zoom lens under constant lighting conditions (including a covering of the microscope to prevent environmental light variation), and using stitch scans of >70 images at the 200× magnification. Images of 1000 x 1000 pixels were then isolated from areas with no damage, including mutant and adjacent wild-type clones from crispant individuals.

All imaging was conducted with constant lighting conditions and acquisition parameters. Whole specimens were imaged using the VH-Z00R lens at a lens magnification set-up of 50x, or a Nikon D5300 camera mounted with a Micro-Nikkor 105mm f/2.8G lens using an f/16 aperture and diffused LED illumination. Microphotographs of wing details were imaged using the VH-Z100T lens at magnification set-ups of 200×, 500×, and 700×. Microphotographs of single scales were acquired using a Nikon D5300 camera mounted on a a Varimag II camera adapter at a 2× magnification setting, a trinocular AmScope ME580-2L metallurgical microscope with a polarizer and analyzer, and a BoliOptics LMPlan Achromatic 50×/NA 0.6 objective.

#### CRISPR mutagenesis

Cas9-2xNLS recombinant protein stock was dispatched in 2.5 μL aliquots after dilution to a concentration of 1000 ng/μL in 2x injection buffer (7.4 pH, 1 mM NaH_2_PO_4_, 1 mM Na_2_HPO_4_, 10 mM KCl). Synthetic sgRNAs were dispatched at 500 ng/μL in 2.5 μL aliquots after dilution in Low EDTA Tris buffer (pH 8.0, 10mM Tris-HCl, 0.1 mM EDTA). Aliquots were stored at −80°C and mixed prior to injection. Eggs were collected from *C. eurytheme* wild-caught gravid females, surface-decontaminated with 5 % Benzalkonium Chloride for 1-2 min, rinsed with distilled water, air-dried and placed upright on double-sided tape rendered less sticky with paper towel fibres. Syncytial embryos up to 4 hrs AEL were micro-injected with a 250:125 ng/μL (for *red*) or 500:250 ng/μL (for *bab*) Cas9:sgRNA mix using pulled borosilicate capillaries. Injection dishes were moved to a plastic container humidity-saturated with a wet paper towel for 48 hrs to avoid desiccation before proper egg healing, and then placed face down on 7-14 d old vetch trays for hatching. All the *bab* crispants characterised in this study were generated in a previous publication (Ficarrotta et al., 2022).

#### UV macrophotography and microphotography

All imaging in the UV-A range was performed using a full-spectrum converted Lumix G3 camera mounted under the illumination of two CFL BlackLight 13 Watt T3 Spiral Light Bulbs (General Electric, USA). All lens set-ups were mounted with a U-Venus-Filter (Baader Planetarium, Germany), which blocks visible light above 400 nm. For imaging of whole specimens, an EL-Nikkor 50 mm f/2.8 N lens was mounted with about 60 mm of extension using focusing helicoids, while for magnified views of wing details, that same lens was reverse-mounted and provided additional extension using a sliding bellows.

#### Scanning Electron Microscopy

Scales of interest were placed on carbon tape and sputter coated with a 10-12 nm layer of gold (Ren et al., 2020). SEM images were acquired on a FEI Teneo LV SEM, using the Everhart-Thornley detector and beam parameters of 2.00 kV / 25 pA with a 10 μs dwell time. Scale cryofracturing was performed as previously described (Ficarrotta et al., 2022; Ren et al., 2020).

#### Image post-processing

No post-processing was performed on the image datasets acquired with the Epson scanner and Keyence microscopes, *ie*. the images used for colour profile analyses. In order to correct for slight overexposure in all visible (Nikon D5300) and UV-A (Lumix G3) images of whole pinned specimens, an overexposure compensation (Black Point value of +30 and White Point value of −15) was applied in post-processing using the Levels function in Adobe Photoshop. This adjustment uniformly changes luminosity and does not affect hues visibly.

### QUANTIFICATION AND STATISTICAL ANALYSIS

#### Automated colour scoring

Images were processed with the *R* package *colordistance* (Weller and Westneat, 2019). Cropped images of each wing and wing surface were imported, and the CIELAB value determined for 10,000 randomly sampled pixels per image. Using the function *getLabHist*, thresholds were applied to remove the white background and brown-black pixels. The remaining pixels within retained *L*a*b\** (Lab) value ranges were placed into 50 bins of the *b\** component (with no subdivisions of *L\** or *a\**), with lower values corresponding to ‘oranger’ wings and high values corresponding to ‘yellower’ wings. This allowed a distribution of pixel hues to be rapidly and repeatedly determined from thousands of images. We took the number of the bin containing the highest number of pixels (the ‘max bin’) value as the phenotypic score for each individual and used this value for QTL analysis. Because this score does not directly correspond to one axis of Lab parameter space, we named it the ‘yellow score’ to reduce ambiguity.

#### Landmarking, morphometrics and clustering analysis

Individual measurements were taken using FIJI (**Figure S7A-B**). Sizes were calibrated with a ruler included in the wing scans. Where possible, pattern elements were measured from left and right wings to ensure consistency of measurements; likewise, size measurements were taken from both the dorsal and ventral wing surfaces. Principal component analysis of wing measurements was performed using the base-R function prcomp(), and plotted with density curves with ggplot2.

Many of the measured variables may be strongly related to each other and therefore contain the same information. In order to identify groups of independent variables we clustered the traits using the R package *ClustOfVar* (Chavent et al., 2012). Briefly, data was log-transformed and a trait hierarchy constructed (**Figure S7D**), and the dispersion of the Rand index for clusters was observed to support the selection of 5 clusters. A latent variable per cluster was then calculated for every individual, and these latent variables were used for the QTL analyses (**Figure S8**).

#### QTL analysis and gene identification

For linkage mapping and QTL analysis, we used the published linkage maps for one F_2_ and two BC broods, which we previously used for genome assembly and mapping of UV iridescent colour (Ficarrotta et al. 2022). For each variable (yellow score and the five cluster latent variables), we performed single QTL genome scans in *R/qtl* (Broman et al., 2003) with Haley-Knott regression, calculating significance thresholds for each scan with 1,000 permutations, and defined 95% Bayes credible intervals for each significant peak.

Two-QTL genome scans were also run with Haley-Knott regression and 500 permutations. The goodness of fit of each significant two-QTL model was examined with the function `fitqtl()` function, which provided the percentage of variance explained by the interaction. We further examined multiple-QTL models, assessing the probability that additional loci might explain a significant fraction of variation when accounting for the detected intervals, though no additional significant QTL were detected this way

