## Supplementary Figures for "Genetics of continuous colour variation in a pair of sympatric sulphur butterflies"

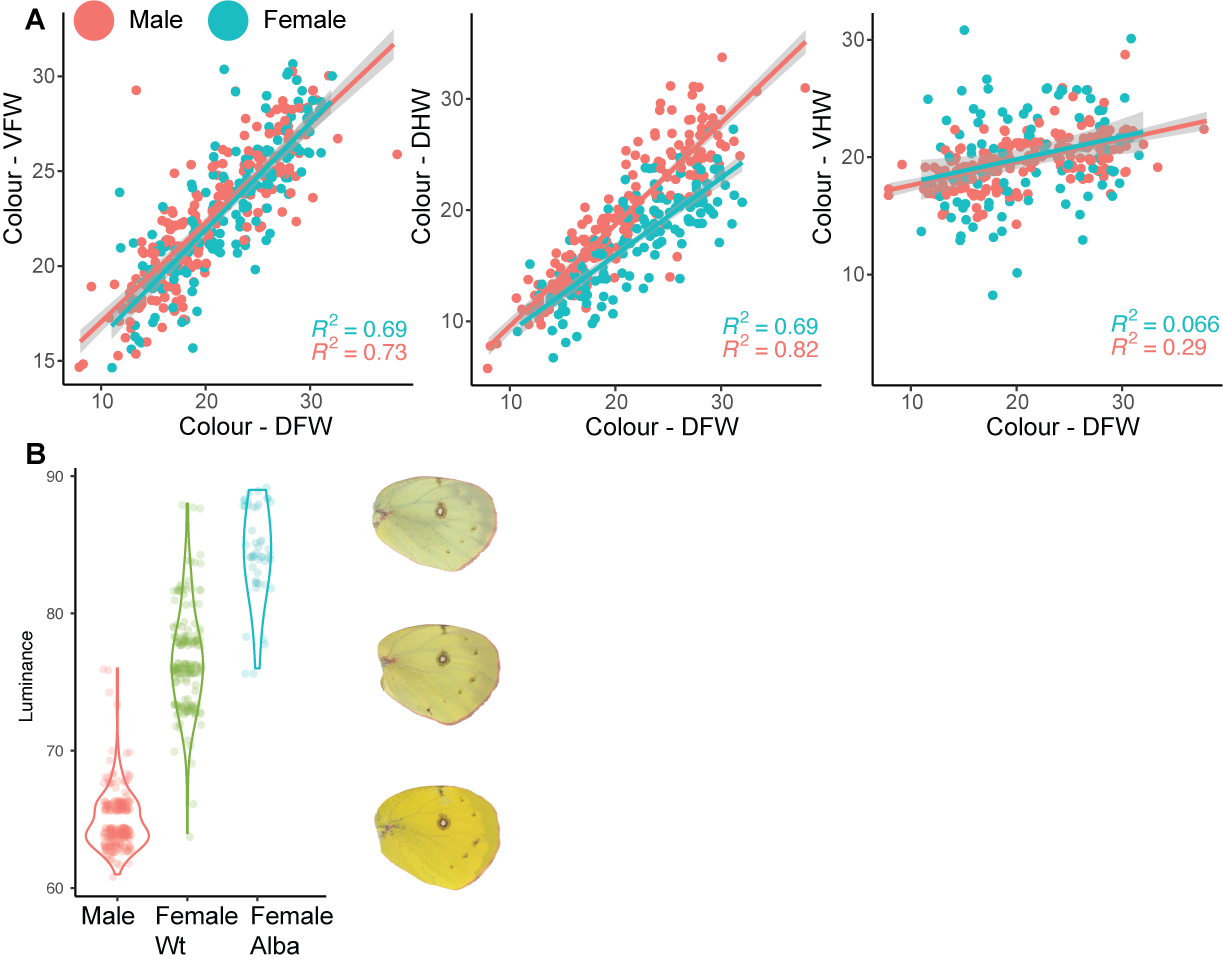

**Figure S1. Correlation of yellow score between wing surfaces of Amherst, MA F_2_ and BC broods used in QTL analysis**. DFW = Dorsal Forewing, DHW = Dorsal Hindwing, VFW = Ventral Forewing, VHW = Ventral Hindwing. **A.** The yellow scores (same values as presented in **Figure 1**) for dorsal forewings are highly correlated with both ventral forewing and dorsal hindwing, in both males and females. However, the yellow scores for the ventral hindwing are not correlated with other wing surfaces and variance is low. **B.** Males and females have variation in luminance values on the ventral hindwing: males have a consistently lower luminance, with females and Alba (unpigmented females) having higher luminance.

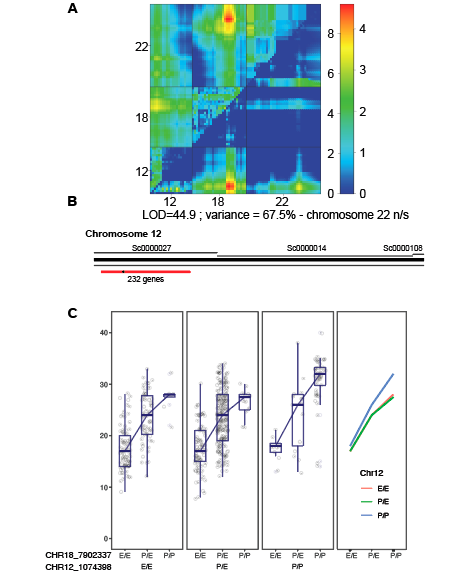

**Figure S2. An additional significant locus for colouration interacts with the chromosome 18 QTL. A**. A two-QTL scan including both backcrosses and the F2 brood detec`ts an additional QTL. A model with the 18, 22 and 12 loci had a LOD score of 44.9 and explained 67.5% of the variance in colour, but indicates that the chromosome 22 locus is not significant. **B**. The LOD interval on chromosome 12 includes 232 genes (supplemental table). **C**. The Phenotype-by-genotype plot for the chromosome 18 and chromosome 12 loci indicates an epistatic interaction; the effect of chromosome 12 variation alone is minimal, but when individuals are homozygous for the *C. philodice* allele at both loci, they are consistently yellower.

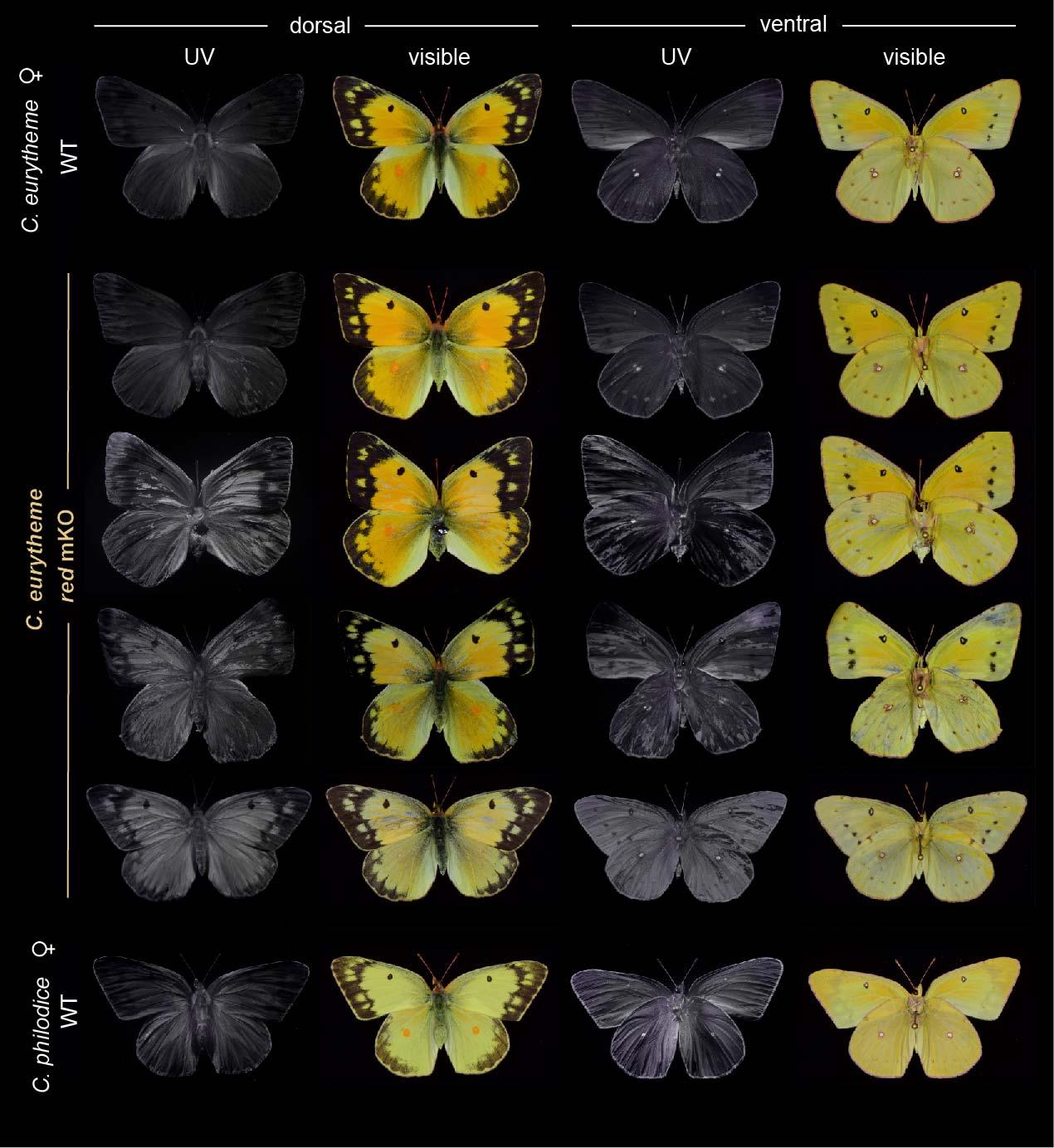

**Figure S3. Representative red crispant phenotypes observed among *C. eurytheme* females.** The extent of crispant mosaics is most visible in the UV-A spectrum, where local loss of pterin pigmentation results in increased UV-reflectance, consistently with the UV-absorbing properties of *Colias* pterin pigments (Wijnen et al. 2007). Ectopic UV-reflectance in pterinless scales is unrelated to the UV-iridescence observed in *C. eurytheme* wild-type males (Figure S4) or in *bab* crispants, which is based on structural modifications of ridge structures on the upper scales (Ficarrotta et al., 2022).

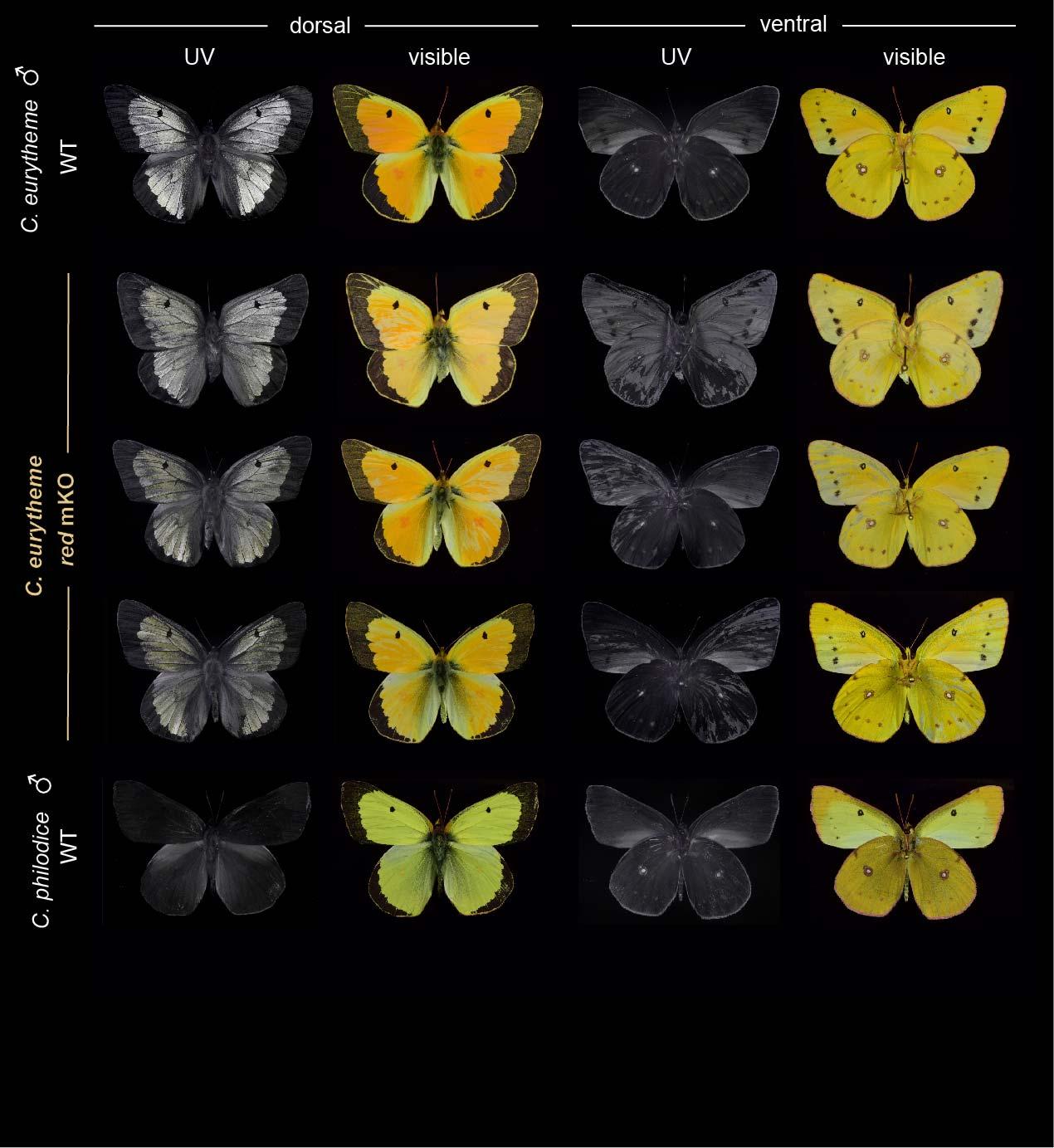

**Figure S4. Representative red crispant phenotypes observed among *C. eurytheme* males.** Effects were comparable with females, with the exception that dorsal effects on UV-reflectance were masked by the strong UV-iridescence of WT *C. eurytheme* males on this wing surface.

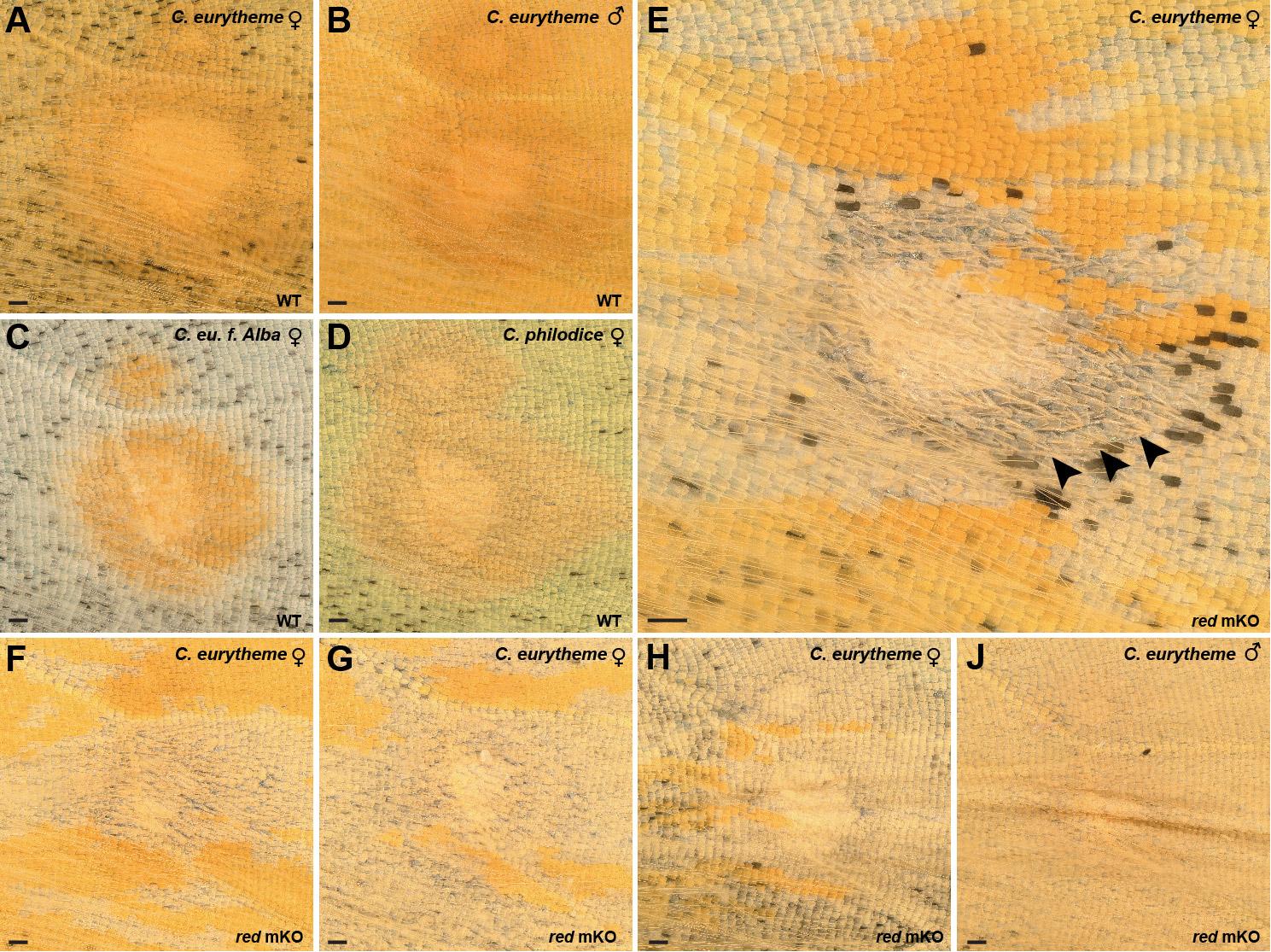

**Figure S5. Magnified views of *red* crispant phenotypes in dorsal hindwing discal spot. A-D**. The discal spot of dorsal hindwings always display a deep orange colour regarding of sex, species, and morph in the *C. eurytheme* / *C. philodice* species pair. **E-J**. Mosaic KOs of *red* consistently result in a loss of orange pigmentation in *C. eurytheme* females. (E-H) and males (J). In rare cases, crispant clones showed defective development, an effect limited to the inner section of the dorsal discal spot, as seen here by the presence of curled scales (arrowheads, panel J). Scale bars : 200 µm.

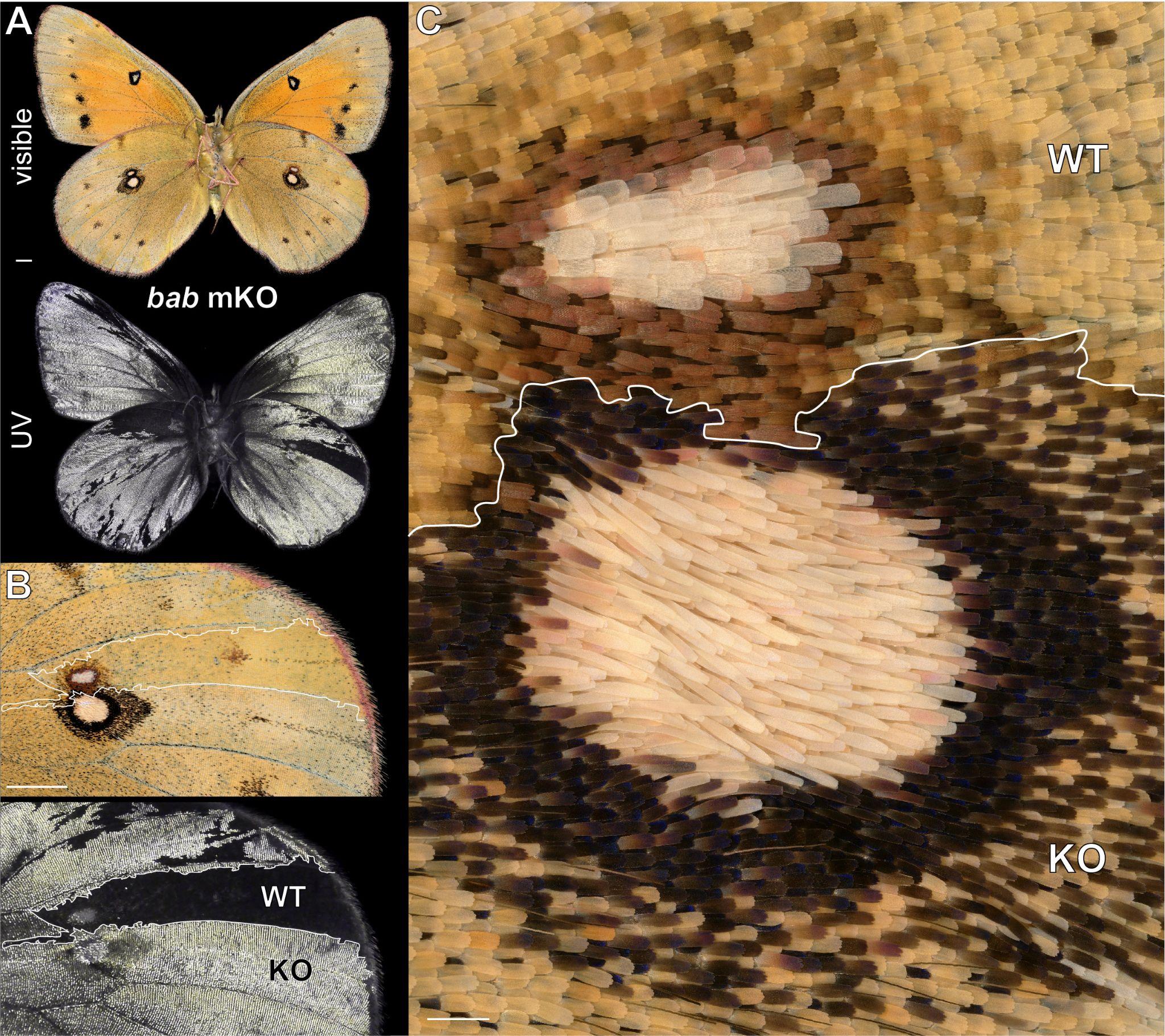

**Figure S6. Pterin-independent effects of *bab* mKOs in ventral hindwing discal spot. A-C**. Ventral wing surface of a female *Colias eurytheme* *bab* crispant with large mutant clones, as seen in the ectopic gain of UV iridescence (bottom panels in A-B). A large *bab*-deficient clone overlaps with the large discal spot in the hindwing and shows scale modifications relative to the adjacent WT spot, including more oblong silver scales, and darker melanic scales in the discal eyespot ring (C). Scale bars : A-B = 2000 µm; C = 200 µm.

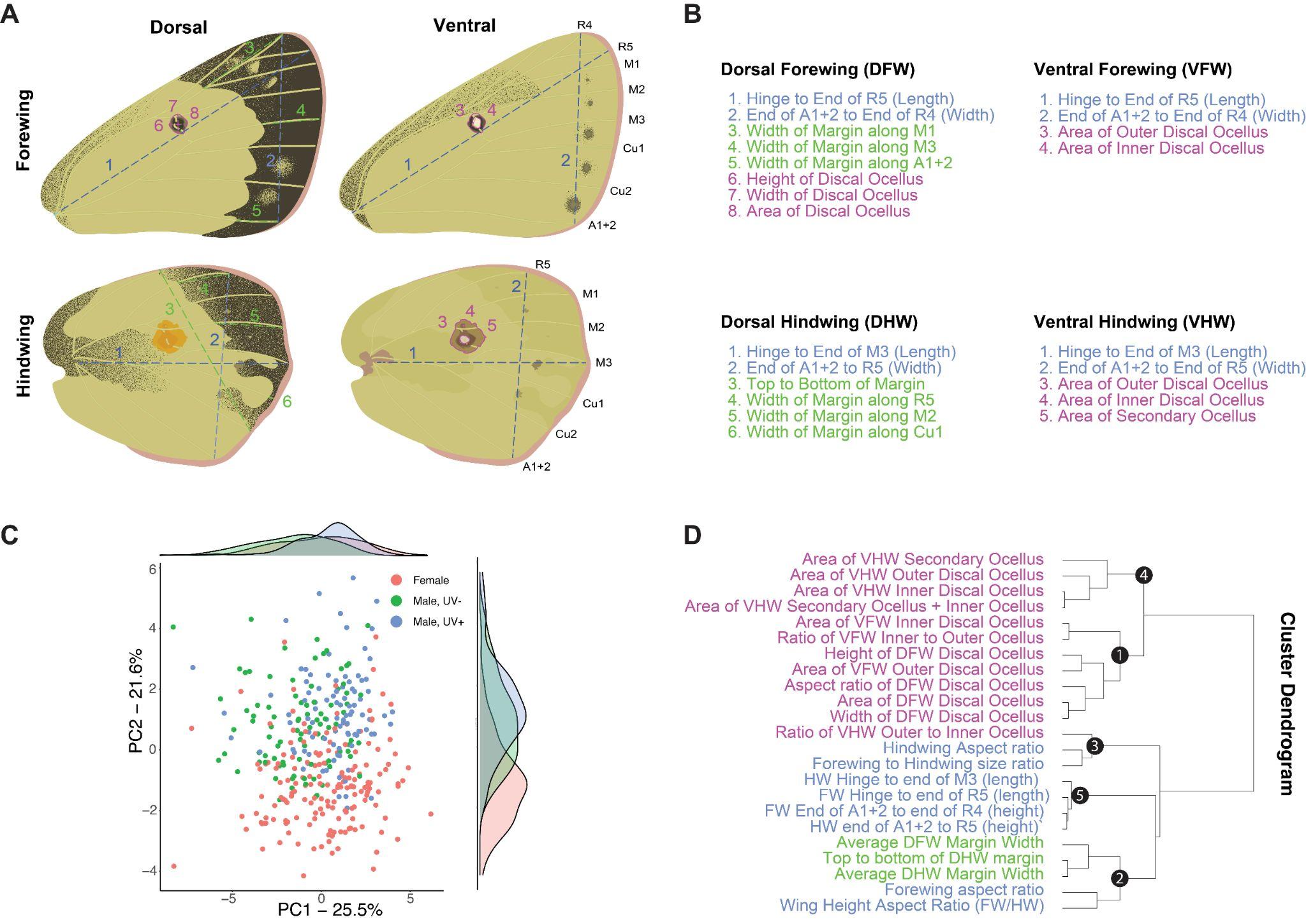

**Figure S7. Morphometric measurement of wing traits used in cluster and QTL analyses. A-B.** Wing size traits (cyan), and wing pattern traits relating to the size of the black wing margin (green) and discal ocelli (magenta), were measured using FIJI on scanned wings from mapping broods. **C.** Principal component analysis of measures, with points coloured by sex and UV status (indicative of Z genotype). UV+ and UV- males are separating on PC1, while males and females are separating on PC2. **D.** Dendrogram of traits and hierarchical clusters (indicated by black nodes). Latent variables for these 5 clusters were used for QTL analysis (**Figure S8**). The Cluster 5 variable ‘*FW hinge to end of M3*’ (forewing length) was used for QTL mapping of wing size (**Figure 7**).

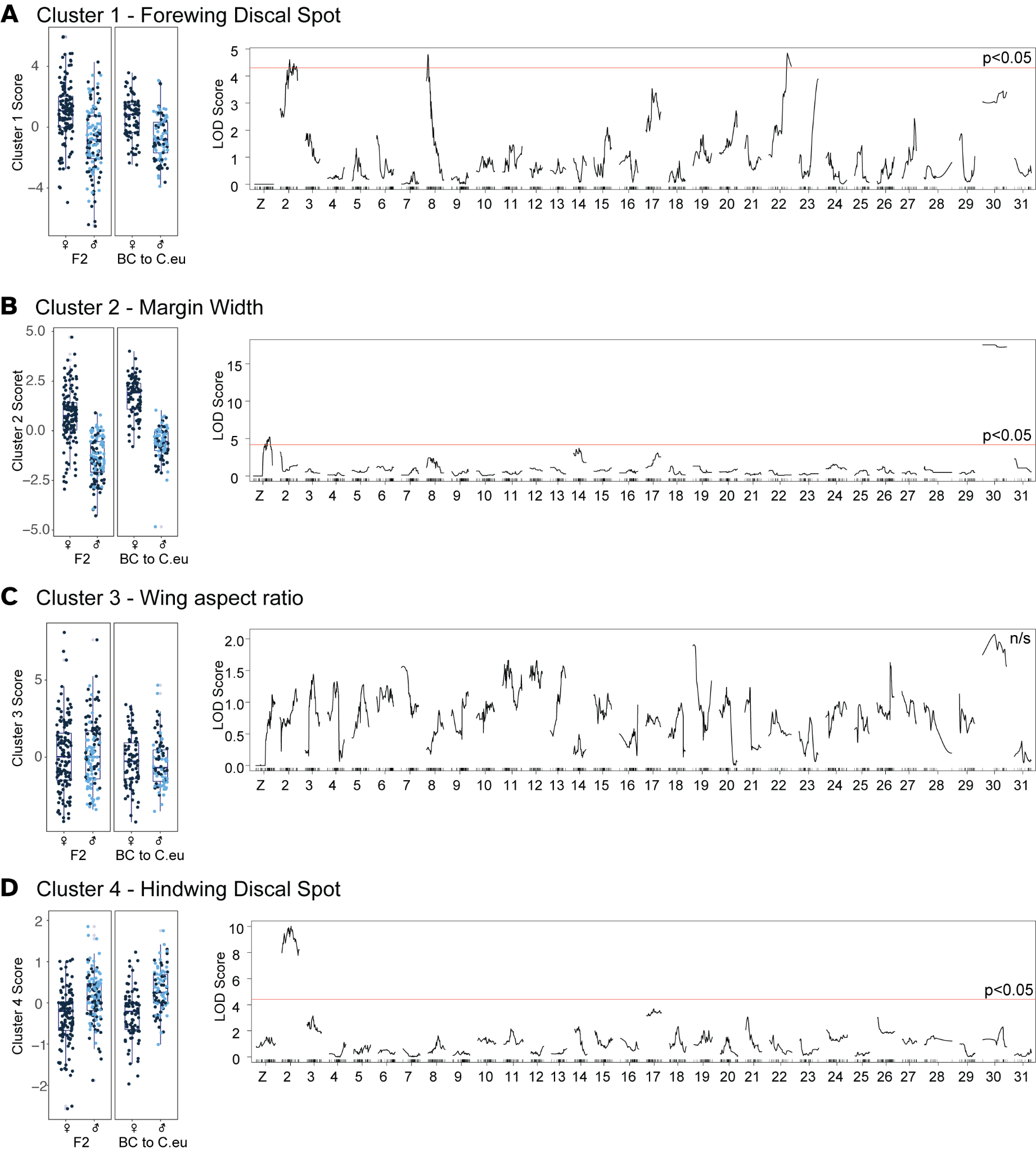

**Figure S8. LOD plots for morphometric trait clusters.** Three significant LOD intervals relating to the size and shape of the discal forewing spot were detected (A), two for the width of the marginal band (B), none for the aspect ratio of the wing (C), and one for the hindwing discal spot (D).

**Colour-enhanced figures**

The main figures of this manuscript are presented below with enhanced hues that are more readily perceived by readers with the most common color vision deficiencies.

Pixels with Hue values 35°-60° (**Figs. 1-2**), or 30°-60° (**Figs. 4-6**), *ie.* the range of orange-yellow pixels observed in the included wing images, were rotated -180° in HSV space using Adobe Photoshop.

Red Channels in RGB space were trasnposed into the Blue Channel to create heatmaps with a magenta-green colour scale using Adobe Photoshop (**Figs. 3 and 7**).

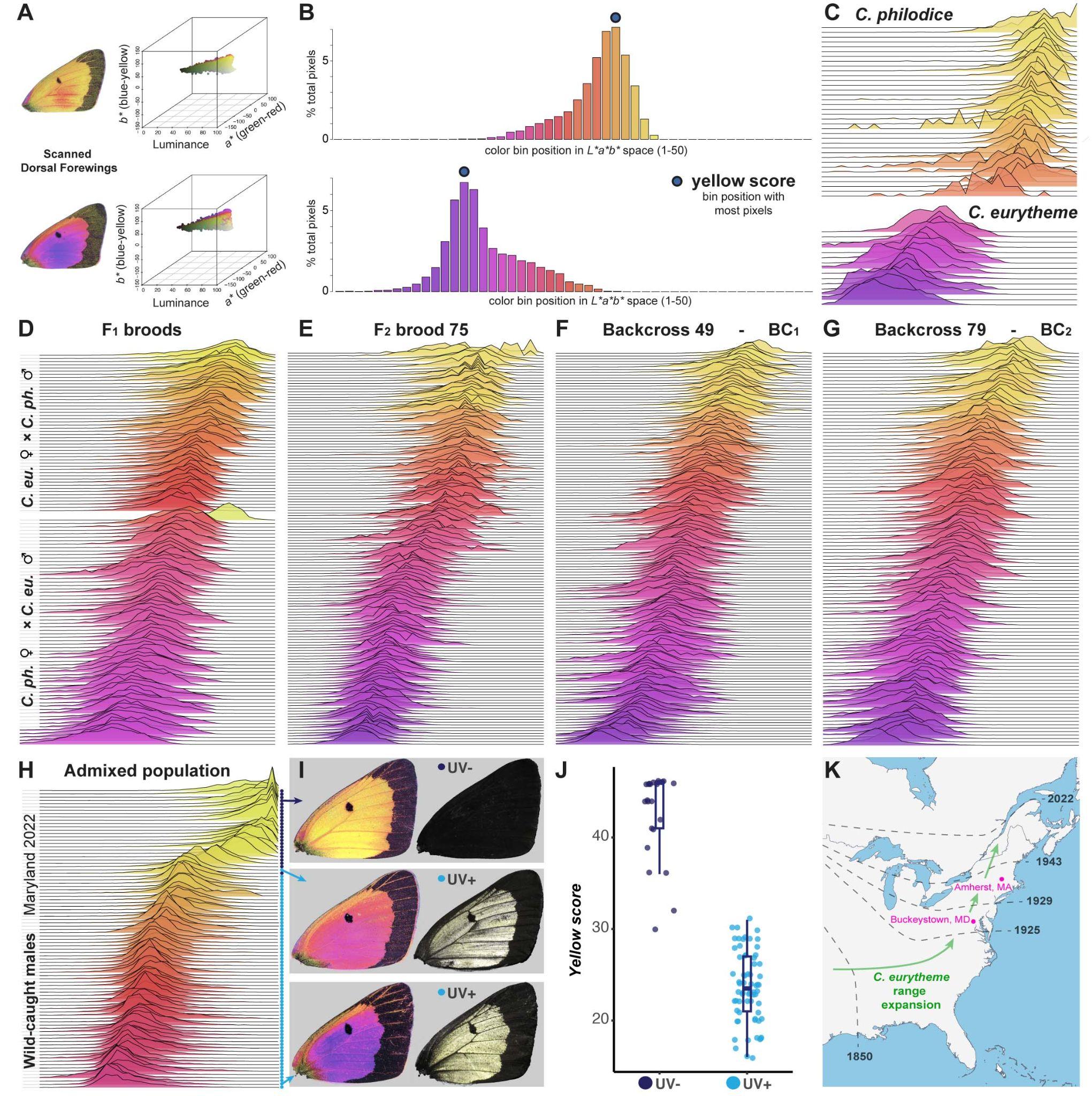
 **Colour-enhanced Figure 1. Rapid, repeatable and automated phenotyping of wing colour**. (**A**) We used the package *r/colordistance* to extract pixel *L*a*b** coordinates from scanned wing images, followed by appropriate thresholding to remove background information. We then selected a subset of the remaining pixel range to capture the orange-yellow region of CIELAB colour space, and divided it into 50 bins along the *b** axis. (**B**) Pixel density histograms for the two individuals shown in A, with colour profile bin positions on the X axis. Bar colours correspond to the average colour of each plotted bin and recapitulate the range of orange-to-yellow pixels of interest in further analyses. The bin with the maximum pixel count is hereafter referred to as ‘yellow score’. (**C-G**) Colour binning profiles of every individual from the offspring of wild-caught individuals (**C**) and indicated broods (**D-G**). (**H-I**) Wild-caught males from an agricultural site where *C. philodice* and *C. eurytheme* fly together and hybridise ; the coloured dots indicate the presence or absence of UV iridescence, allowing inference of the Z chromosome genotype (Ficarrotta et al, 2022). (**J**) UV-positive males are consistently oranger while UV-negative males are consistently yellower. (**K**) Summary of the northward expansion of *C. eurytheme* across the Eastern US after 1850, as previously documented (Hovanitz, 1944). The Eastern US comprised part of the native range of *C. philodice* before 1850. Parental admixed populations used for QTL mapping (C-J) originated from Amherst, MA (Summer 2000). Wild-caught males (H-I) were collected in Buckeystown, MD (Aug-Sep 2022).

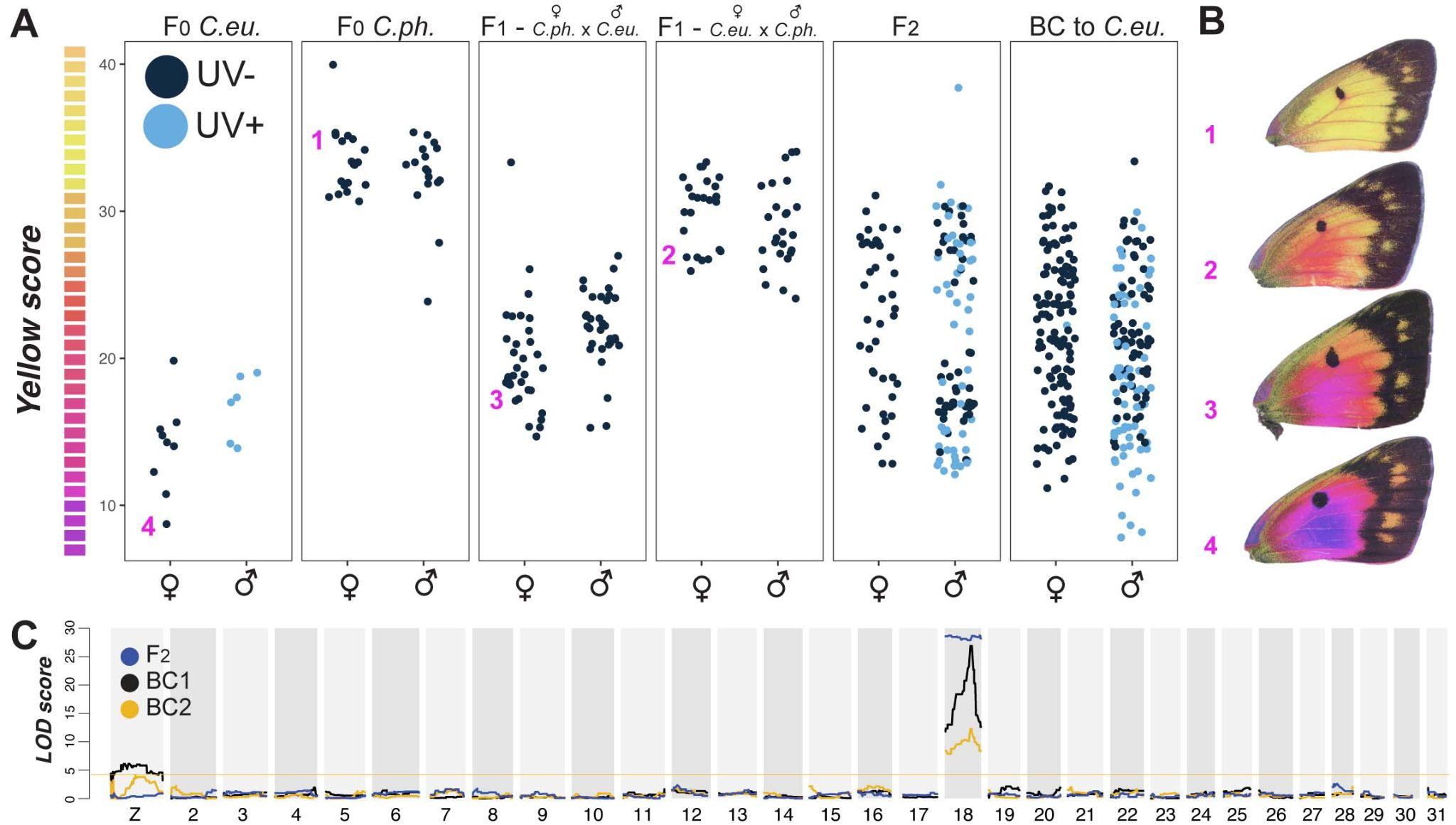

**Colour-enhanced Figure 2. Yellow-orange colour variation is linked to chromosome 18 and the Z sex chromosome**. We measured yellow scores for all individuals from one F_2_ and two backcross broods, as well as for a random subset of individuals from grandparental F_0_ and parental F_1_ broods. F_1_ and F_2_ broods show intermediate colour between the parental broods, (**A**). Four exemplar individuals from different parts of the colour range are depicted in (**B**). We used these yellow scores as the input into QTL analyses, and found a LOD interval on chromosome 18 for all three broods, as well as LOD interval on the Z chromosome in both backcross broods (**C**).

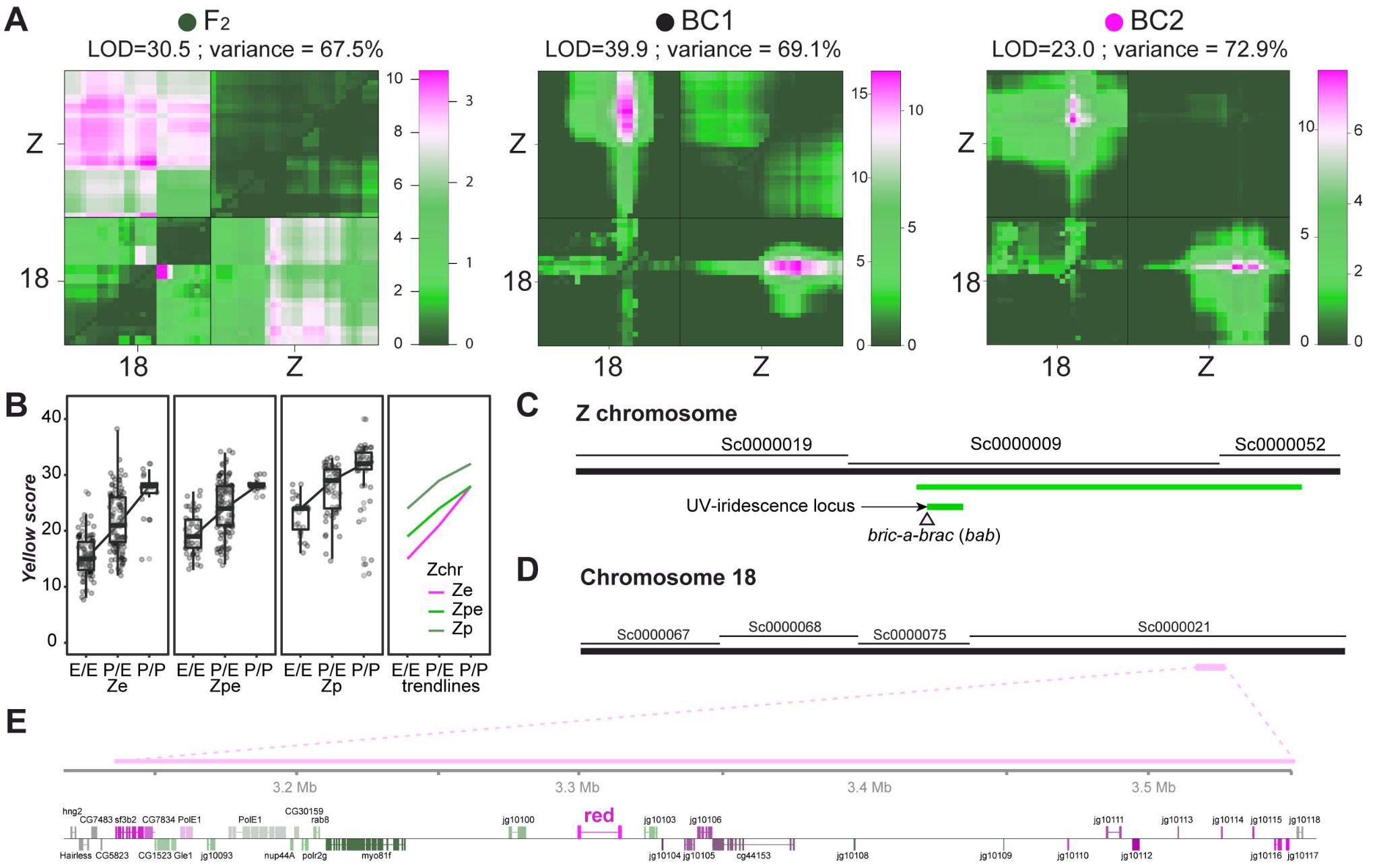

**Colour-modified Figure 3. Additive interaction between the LOD intervals on chromosome 18 and the Z chromosome. A.** Two-QTL models for each brood indicate a significant interaction between the Z chromosome and chromosome 18. For each plot, the top left corner is the full model (two-QTL model plus interaction) and tests for epistasis, whereas the lower right tests for additivity among loci (two-QTL without interaction). Each colour scale indicates two separate LOD scores for epistasis (left) and additivity (right). Significance threshold is set at 4.7 based on previous recommendations (Broman et al., 2003). **B.** Genotype x Phenotype plots showing the relationship between the Z chromosome genotypes (*Ze* and *Zp*: homozygotes or hemizygotes; *Zpe*: heterozygotes) and chromosome 18 LOD interval genotype (x-axis) on yellow score. The rightmost panel overlays the means for the first three panels. **C.** The Z chromosome includes the U locus (cyan) locus for UV-iridescence (Ficarrotta et al., 2022). **D-E**. The LOD interval on Chromosome 18 (yellow) is centred on the gene *red Malpighian tubules* (*abbr.* *red*).

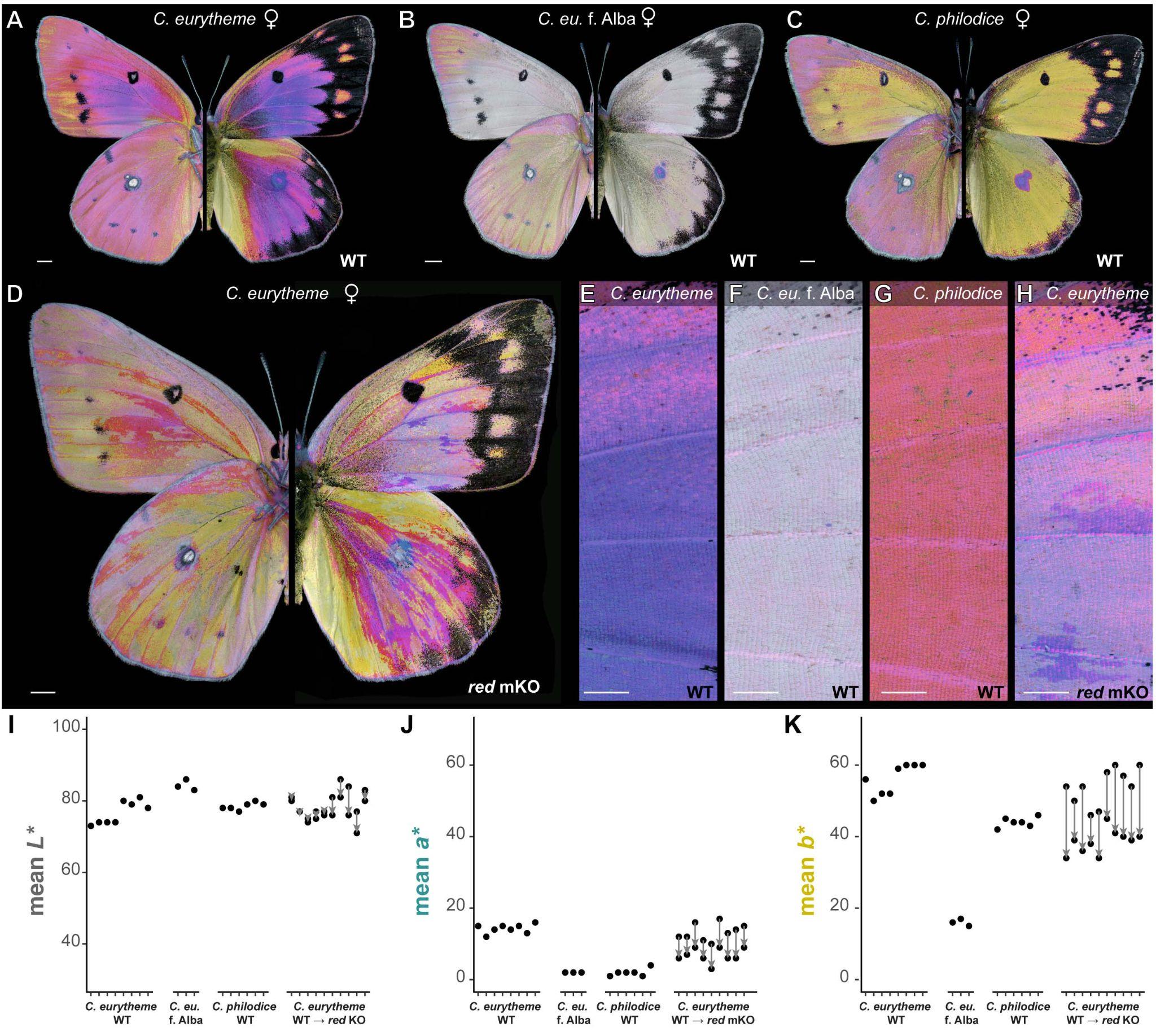

**Colour-Enhanced Figure 4. Wing colouration effects of *red Malpighian tubules* CRISPR mKOs. A-C.** Ventral (left) and dorsal of wild-type (WT) females representative of admixed *C. eurytheme* and *C. philodice* populations, here sampled from Maryland. **D.** *C. eurytheme* G_0_ mosaic crispant for the *red* gene with discoloured streaks (mutant clones), imaged in identical conditions to the wild-type controls. **E-H**. Side-by-side comparisons of dorsal forewing regions show reduced orange colouration in *red* KO clones (panel H) relative to *C. eurytheme* WT clones. **I-K**. Comparison of mean *L** (luminance), *a**(green to red), and *b** (blue to yellow) pixel values sampled from dorsal WT forewings, and from adjacent WT vs. *red*-deficient clones from 10 *C. eurytheme* crispants. Scale bars : A-D = 2,000 µm ; E-H = 1,000 µm.

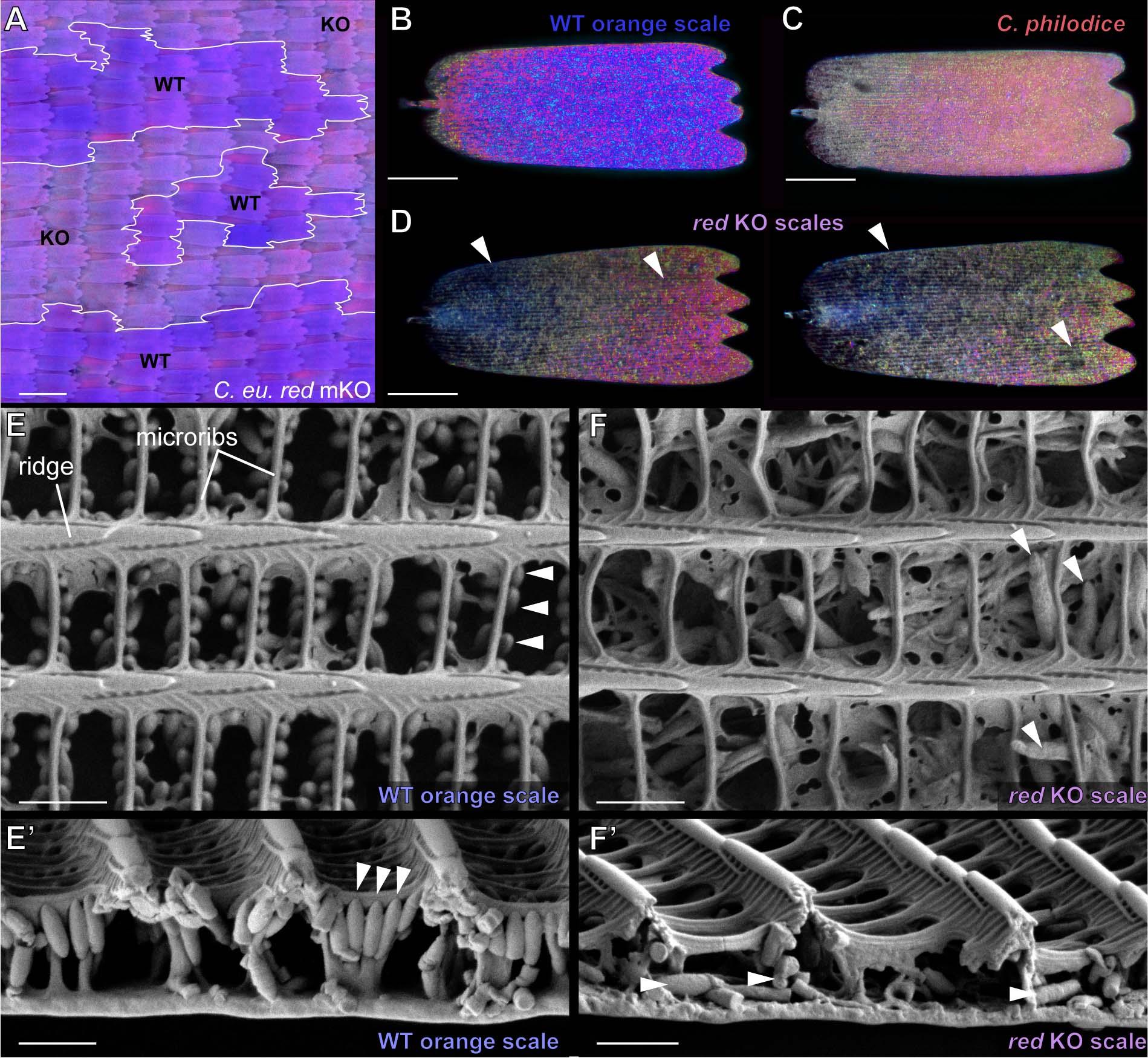

**Colour-enhanced Figure 5. Disorganisation of internal scale structures in *red* crispant scales**.

**A.** Scale-level colouration of *red* mKO clones relative to wild-type (WT) phenotypes in *C. eurytheme*, here in dorsal forewing regions. **B-D**. Reflected polarised microscopy of individual dorsal forewing scales from a *C. eurytheme* female (**B**), a *C. philodice* female (**C**), and a female *C. eurytheme* *red* crispant clone (**D**). Scales from panels B and D are from adjacent clones in the same crispant individual. Scales deficient for *red* have poor refringence and show an overall lack of pigment density (arrowheads). **E-E’**. SEM top (**E)** and cross-sectional (**E’**) views of WT orange scales. Pterin granules (arrowheads) are visible as oblong structures attached to the microribs, the structures that form transversal bridges between the ridges of the scale upper lamina. **F-F’**. Defects of the inner scale lumen in *red* crispant scales, amidst a normal upper surface. Pterin granules (arrowheads) are improperly formed, fail to attach to microribs, and are randomly arranged in a disorganised scale matrix. Scale bars : A = 100 µm ; B-D = 50 µm ; E-F’ = 1 µm.

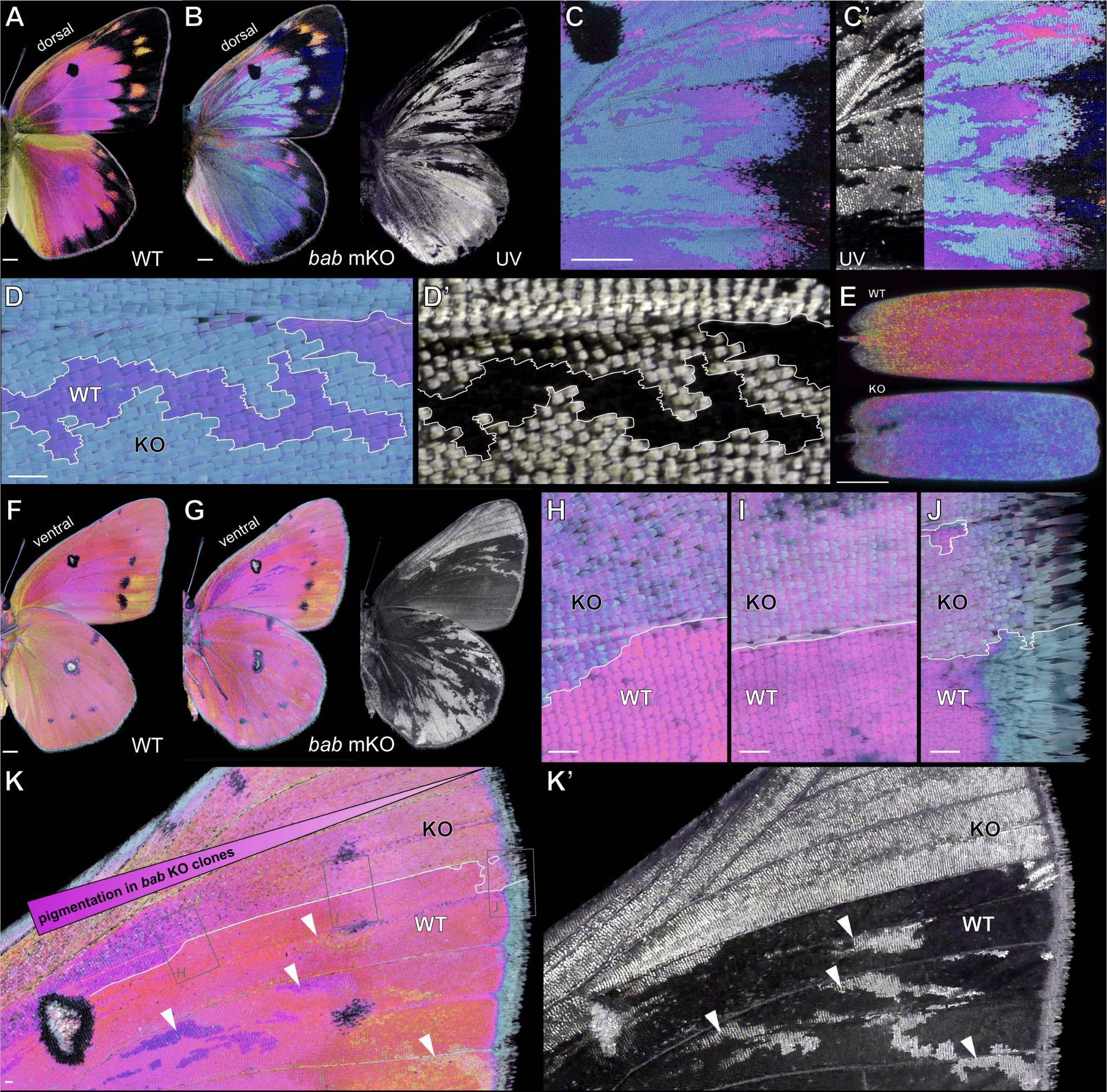

**Colour-enhanced Figure 6. CRISPR mutagenesis of *bab* can affect both UV-iridescence and pterin pigmentation. A.** WT *C. eurytheme* female, dorsal view. **B-C’**. Dorsal views of a *bab* crispant *C. eurytheme* female (Ficarrotta et al., 2022). Crispant clones lacking *bab* display ectopic UV-iridescence in all regions, ectopic blue iridescence over melanic regions, and colour shifts in orange regions, with the two latter effects only observed in crispant females. C shows the visible-spectrum colouration with normal light incidence (0°), while C’ reveals the structural iridescence effects in the UV-A and visible ranges observed with a 30° light incidence. **D-D’.** Magnified views of WT and KO *bab* crispant clones with 0° incidence. **E**. Polarised reflective microscopy of adjacent scales from across a *bab* crispant boundary. **F**. WT *C. eurytheme* female, ventral view. **G-K’**. Ventral views of a *bab* crispant *C. eurytheme* female. On the ventral surface, the effect of *bab* loss-of-function over pterin pigmentation changes over the proximo-distal axis. Scale bars : A-B, F-G = 2,000 µm ; C-D, H-K = 200 µm ; E = 50 µm.

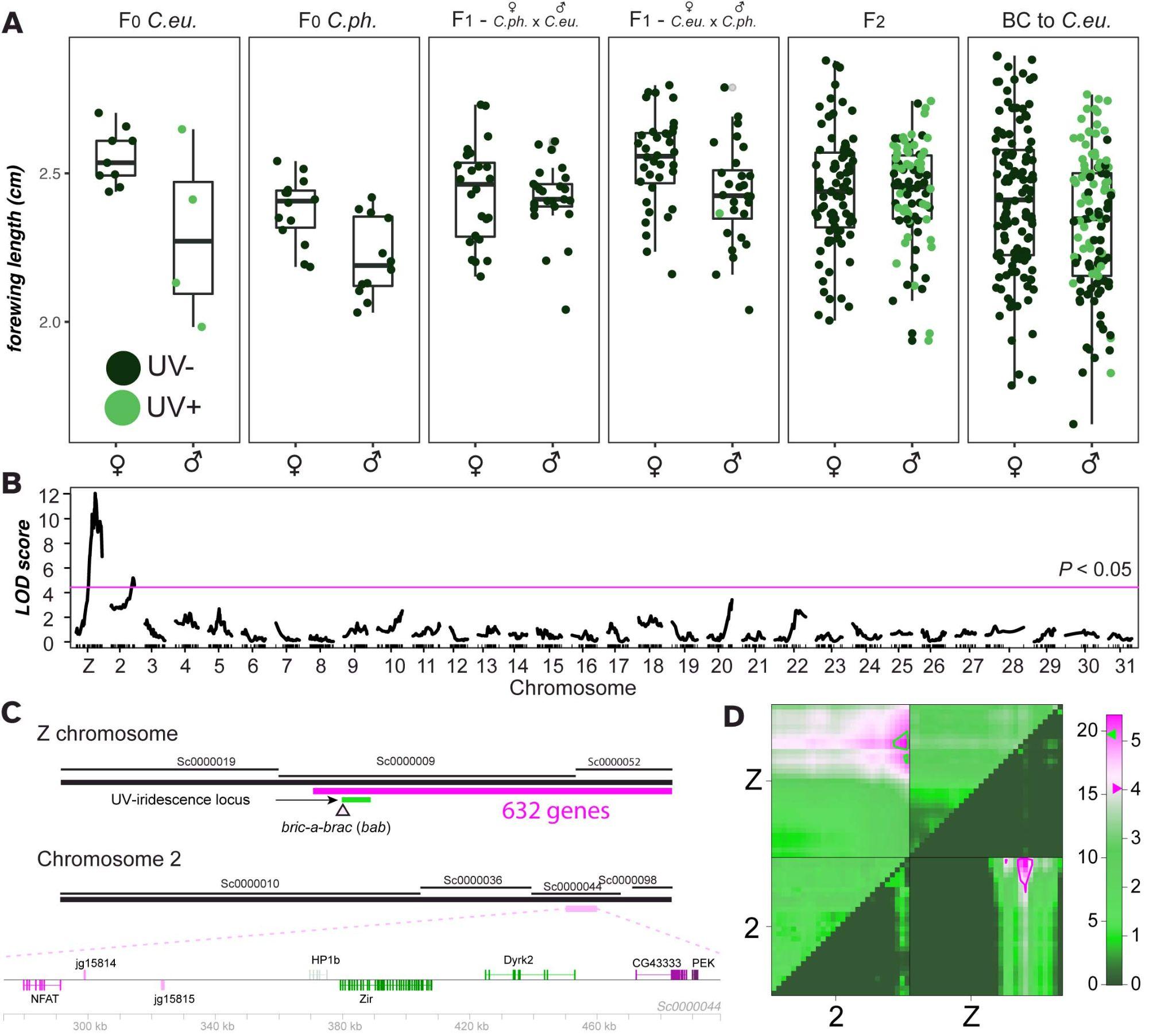

**Colour-modified Figure 7. Two LOD intervals associated with wing size. A.** Variation in wing length in crosses. **B-C.** A single-QTL model detects two significant loci (B), which incorporate large portions of chromosome 2 and the Z chromosome (C). **D.** Support for an additive two-QTL model that explains 18% of the variation in size, with LOD scores for the full model on the upper left and for the additive model on the lower right.
